# A novel AR molecular glue degrader via bivalent engagement of the N-terminal domain and RNF213–UBE2J2 complex

**DOI:** 10.64898/2026.07.14.738453

**Authors:** Kunzhong Wu, Qiong Wu, Yongzhi Lu, Qi Yang, Ting Ran, Jiyun Zhang, Xianjie Qiu, Chengjie Huang, Yue Lin, Jiayi Chen, Zelin Yang, Jie Zhang, Wei Qin, Zhu Liu, Xinqi Liu, Miru Tang, Hongming Chen, Jie Zheng, Xinwen Chen, Jinsai Shang

## Abstract

Targeted protein degradation enables the elimination of disease-relevant proteins through induced proximity between E3 ligases and substrates. Current androgen receptor (AR) therapies act via the ligand-binding domain (LBD) but fail against constitutively active LBD mutants or splice variants such as AR-V7. Here, we identify GZL626, a small-molecule degrader that induces proteasome-dependent degradation of both full-length AR and AR-V7. GZL626 suppresses AR transcriptional activity, inhibits proliferation of AR-positive prostate cancer cells, and blocks SARS-CoV-2 infection. Mechanistically, GZL626 binds the intrinsically disordered AR N-terminal domain (NTD) and promotes formation of a noncanonical RNF213-UBE2J2-AR ternary complex that drives AR ubiquitination and degradation. The AR DNA-binding domain (DBD) acts as a structural hub within this assembly. Distinct from PROTACs or classical molecular glues, GZL626 engages both AR and RNF213 bivalently, remodeling protein interactions to stabilize a functional E3-E2-substrate complex. These findings establish GZL626 as a first-in-class NTD degrader with potential to overcome resistance in advanced prostate cancer.

## Introduction

The androgen receptor (AR) is a ligand-activated transcription factor of the nuclear receptor superfamily that plays pivotal roles in modulating gene networks essential for sexual development, metabolic homeostasis, and immune regulation^1^. AR comprises three functional domains: a conserved DNA-binding domain (DBD) that recognizes androgen response elements, a ligand-binding domain (LBD) that confers specificity for steroid hormones such as testosterone and dihydrotestosterone (DHT), and a highly flexible N-terminal domain (NTD) that mediates transcriptional activation through interactions with coregulators and chromatin-remodeling machinery^2–4^. Dysregulation of AR signaling drives some pathologies, most prominently prostate cancer and androgenetic alopecia, which highlights its broad therapeutic significance.

Prostate cancer is one of the most prevalent malignancies in men worldwide^5^. Although localized disease can often be cured, advanced and metastatic prostate cancer depends largely on androgen deprivation therapy, typically combined with AR antagonists^6^. Although the cure is initially effective, nearly all patients relapse and progress to castration-resistant prostate cancer (CRPC), characterized by persistent AR signaling sustained by gene amplification, point mutations, or expression of constitutively active splice variants such as AR-V7^7,8^. Because AR-V7 lacks the LBD, it is resistant to therapeutic agents and remains transcriptionally active^9^, yet no AR-NTD inhibitor or degrader has been applied to clinical treatment. Early candidates, including EPI-family inhibitors and UT-family degraders, performed limited potency or poor pharmacokinetics, while next-generation compounds remain hampered by weak structural scaffolds, incomplete SAR insights, and the structural disorder of the NTD itself ^9–14^.

Beyond oncology, AR has attracted attention in infectious diseases. During the SARS-CoV-2 pandemic, epidemiological data suggested that men were at higher risk of severe COVID-19 and the ADT conferred partial protection^15^. TMPRSS2, a major protease in SARS-CoV-2 entry, is an androgen-regulated gene^16^. Several FDA-approved AR antagonists (enzalutamide, apalutamide, darolutamide) and clinical-stage AR degraders (ARV-110, ARD-61, proxalutamide) downregulates TMPRSS2 and ACE2 expression, inhibiting SARS-CoV-2 infection in prostate-derived and lung cell models.^16–18^. Notably, proxalutamide showed broad-spectrum activity against viral variants and synergized with remdesivir^18^. However, TMPRSS2 regulated by AR is tissue-specific, and enzalutamide failed to show antiviral effects in lung epithelial cells^19^. Large clinical trials, including COVIDENZA (NCT04475601) and the Hormonal Intervention (NCT04397718), subsequently demonstrated that enzalutamide or ADT was not beneficial to hospitalized patients, and concerns over trial design led to retraction of early proxalutamide reports^18,20,21^. The contradictory findings have cast doubt on the utility of AR antagonists as antiviral agents and highlight the need for novel strategies that can overcome the limitations of conventional LBD-targeted drugs.

Targeted protein degradation (TPD) has emerged as a transformative approach to eliminate disease-causing proteins previously considered undruggable^22^. Both proteolysis-targeting chimeras (PROTACs) and molecular glue degraders (MGDs) exploit the ubiquitin-proteasome system to induce selective target degradation^23^. PROTACs employ a modular architecture that links E3 ligase ligands (typically CRBN/VHL) to target binders, offering versatility but often resulting in high molecular weight and suboptimal pharmacokinetic^22,24^. In contrast. MGDs are typically small, monovalent molecules (<500 Da) that reprogram protein-protein interactions to trigger degradation^25–27^, but their discovery has largely been serendipitous due to the complexity of neo-surface formation and the absence of predictive design frameworks^22,23^. Despite these challenges, both modalities have delivered clinical proof-of-concept against key oncogenic proteins, including AR, ER, EGFR, and KRAS^23,28–30^.

Here, we discovered GZL626, a dual-function degrader, targets both AR and the splice variant AR-V7. GZL626 potently suppresses the proliferation of hAR-expressing cell lines and inhibits SARS-CoV-2 infection. Mechanistically, GZL626 acts as a bivalent molecular glue that engages RNF213 and AR simultaneously, promoting UBE2J2 recruitment and formation of a ternary E3-AR-E2 degradation complex. This unique modality enables efficient AR ubiquitination and proteasomal elimination. By overcoming the limitations of classical antagonists and expanding the therapeutic scope of AR degradation, GZL626 provides a powerful tool for targeting resistant prostate cancer and exploring host-directed antiviral strategies.

## Results

### Identification of dual-function AR and AR-V7 degrader

To overcome the limitations of LBD-targeting antagonists in CRPC^14,29,31^, we performed a compound library screen using AR transactivation and 22Rv1 proliferation assays, with ARV110 as a positive control. 9 hits that exhibited potent hAR-antagonism and inhibited 22Rv1 proliferation without affecting AR-ΔNTD transactivation, which suggests their activity was achieved through the NTD (**Fig. 1a, b, and Extended Data Fig. 1a-d**). Among the hits, GZL625, GZL626, GZL627, and GZL628 efficiently reduced both AR and AR-V7 expression levels in 22Rv1 cells at 10 μM, with GZL626 emerging as the most potent (**Fig. 1c**). GZL626 induced time- and dose-dependent degradation of both isoforms, which was blocked by the proteasome inhibitor carfilzomib, whereas the neddylation inhibitor MLN4924 exhibited a modest stabilizing effect, suggesting that GZL626 primarily engages a proteasome-dependent, CRL-independent degradation pathway. (**Fig. 1d and Extended Data Fig. 1f, g**). As expected, the AR degrader ARV110 effectively degraded full-length AR (DC_50_ = 8.6 nM) but did not affect AR-V7 (**Fig. 1e and Extended Data Fig. 1e**). In contrast, GZL626 potently degraded both AR and AR-V7 with comparable potency to EN1441 and no detectable activity against PPARγ, ER, or GR (**Fig. 1e**). Degradation was also observed in HEK293T and LNCaP cells (**Extended Data Fig. 1h-j**), with a DC_50_ of 6.0 μM determined by AR-based AlphaLISA (**Fig.1f**). Given that clinically relevant AR LBD mutations such as H875Y and F877L frequently emerge in CRPC, we further tested whether GZL626 could degrade these mutants. In AR-knockout HEK293T cells overexpressing AR H875Y or F877L, GZL626 efficiently reduced both mutant protein levels (**Extended Data Fig. 1k, l**), demonstrating that its activity extends to other clinically common LBD mutants. These findings confirm that GZL626 selectively targets both AR (including wild-type, T878A, H875Y, F877L) and AR-V7 without detectable off-target effects on other nuclear receptors.

**Fig. 1.**
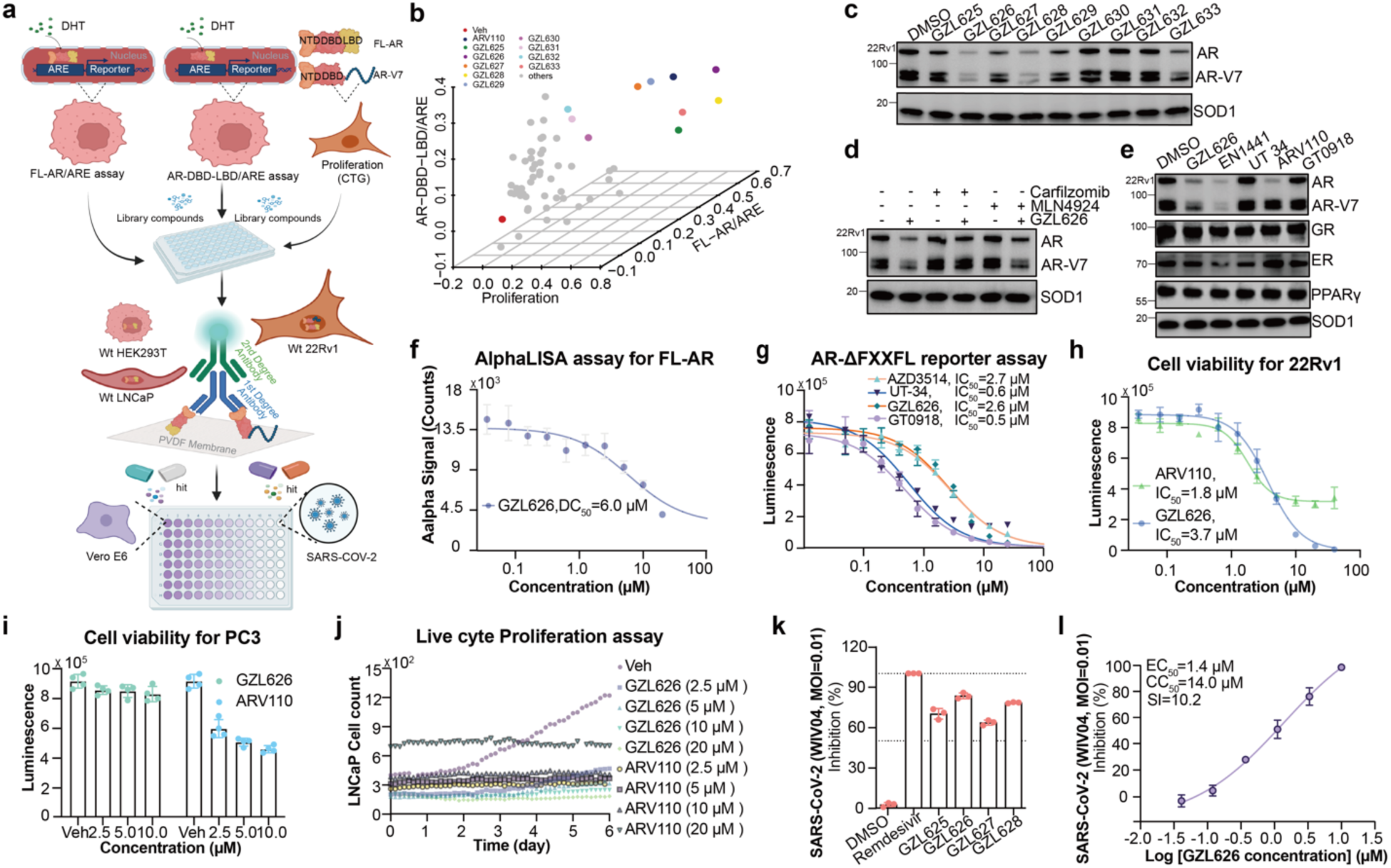
Identification of dual-function AR and AR-V7 degrader. **a,** Schematic of biochemical screening assays used to identify dual-function AR/AR-V7 degraders. **b,** 3D scatter plot showing compound library effects on AR transactivation (full-length AR and AR-ΔNTD) and 22Rv1 cell proliferation. **c,** AR and AR-V7 protein levels in 22Rv1 cells treated with vehicle (DMSO) or GZL626 and its derivatives (10 μM, 24 h), detected by western blotting. **d,** AR and AR-V7 levels in 22Rv1 cells pretreated with GZL626 (10 μM, 4 h), followed by vehicle, carfilzomib (10 μM), or MLN4924 (10 μM) for 8 h. **e**, GZL626 selectively degrades AR in 22Rv1 cells. Protein levels of AR, GR, ER, and PPARγ were assessed following treatment with GZL626 versus EN1441, UT-34, ARV110, and GT0918 under identical conditions (10 μM, 12 h). **f,** Dose-dependent AR degradation in LNCaP cells treated with vehicle or GZL626 (20-0.036 μM, 2-fold serial dilutions, 24 h), measured with an AR total detection kit. **g,** Inhibition of AR-ΔFXXFL reporter activity by GZL626 in the presence of 20 nM DHT; UT-34 and AZD3514 were included as positive controls. **h**-**i,** Growth inhibition of 22Rv1 (**h**) and PC3 (**i**) cells by GZL626 or ARV110 (20-0.036 μM, 2-fold serial dilutions), with calculated IC_50_ values. **j,** Proliferation of LNCaP cells treated with vehicle, GZL626 (20-2.5 μM, 2-fold serial dilutions), or ARV110 as control, monitored in real time for 6 days using a label-free high-content system. **k,** Inhibition of SARS-CoV-2 infection in Vero E6 cells by GZL626, its derivatives, or remdesivir. **l,** Dose-dependent inhibition of SARS-CoV-2 infection by GZL626 in Vero E6 cells, with calculated EC_50_ and CC_50_ values. **b, f-l,** n = 3 independent experiments. **f-l,** data shown as mean ± s.d. All western blot data are representative of three independent measurements

Functionally, GZL626 suppressed AR-Δ-FXXFL transactivation with an IC_50_ of 2.6 μM in the presence of 20 nM DHT, comparable with AZD3514 (2.7 μM), GT0918 (0.5 μM), and UT-34 (0.6 μM) (**Fig. 1g**)^3^. Deletion of the FXXFL motif, which mediates AR N-C interactions, did not affect antagonism, suggesting a distinct NTD-based mechanism. Consistently, GZL626 selectively inhibited proliferation of AR-positive prostate cancer cells, with IC_50_ values of 3.7 μM in 22Rv1 and 4.0 μM in LNCaP cells, while showing minimal effects in AR-negative PC3 cells. In contrast, ARV110 inhibited both AR-positive and AR-negative cells, consistent with off-target toxicity (**Fig. 1h, i and Extended Data Fig. 2a, b**). We further assessed the effects of GZL626 and ARV110 on proliferation and random motility of LNCaP cells using a label-free high-content system. GZL626 was comparable to ARV110 at the same concentration in suppressing cell number, total dry mass, and migration speed (**Fig. 1j and Extended Data Fig. 2c, d**). These findings suggest that GZL626 activity is dependent on AR expression, whereas ARV110 exhibits AR-independent cytotoxicity^32^. Given the role of AR in regulating TMPRSS2, we also tested the compound set (**Extended Data Fig. 3**) in Vero E6 cell-based SARS-CoV-2 infection CPE assay. GZL626 and GZL628 inhibited viral infection, with IC_50_ values of 1.4 μM and 1.8 μM, respectively, showing efficacy comparable to remdesivir at 5 μM (**Fig. 1k, l and Extended Data Fig. 2e**). Taken together, these results identify GZL626 as a dual-function degrader of AR and AR-V7 that suppresses AR transcriptional activity, selectively inhibits AR-driven prostate cancer cell growth, and exhibits antiviral activity against SARS-CoV-2.

### GZL626 suppresses AR-dependent transcription and downstream signaling

To define GZL626 induced global effects, we performed transcriptomic and proteomic profiling in androgen-dependent LNCaP cells treated with GZL626 or vehicle. Integrated analysis revealed selective downregulation of AR target genes including *KLK3* and *NKX3-1*, along with enrichment of pathways linked to apoptosis, cell cycle arrest, and suppression of prostate cancer and viral infection (**Fig. 2a, b and Extended Data Fig. 4a-d**). Consistently, qRT-PCR confirmed significant reductions in *KLK2*, *KLK3,* and *TMPRSS2* in both LNCaP and 22Rv1 cells after GZL626 treatment (**Fig. 2c**).

**Fig. 2.**
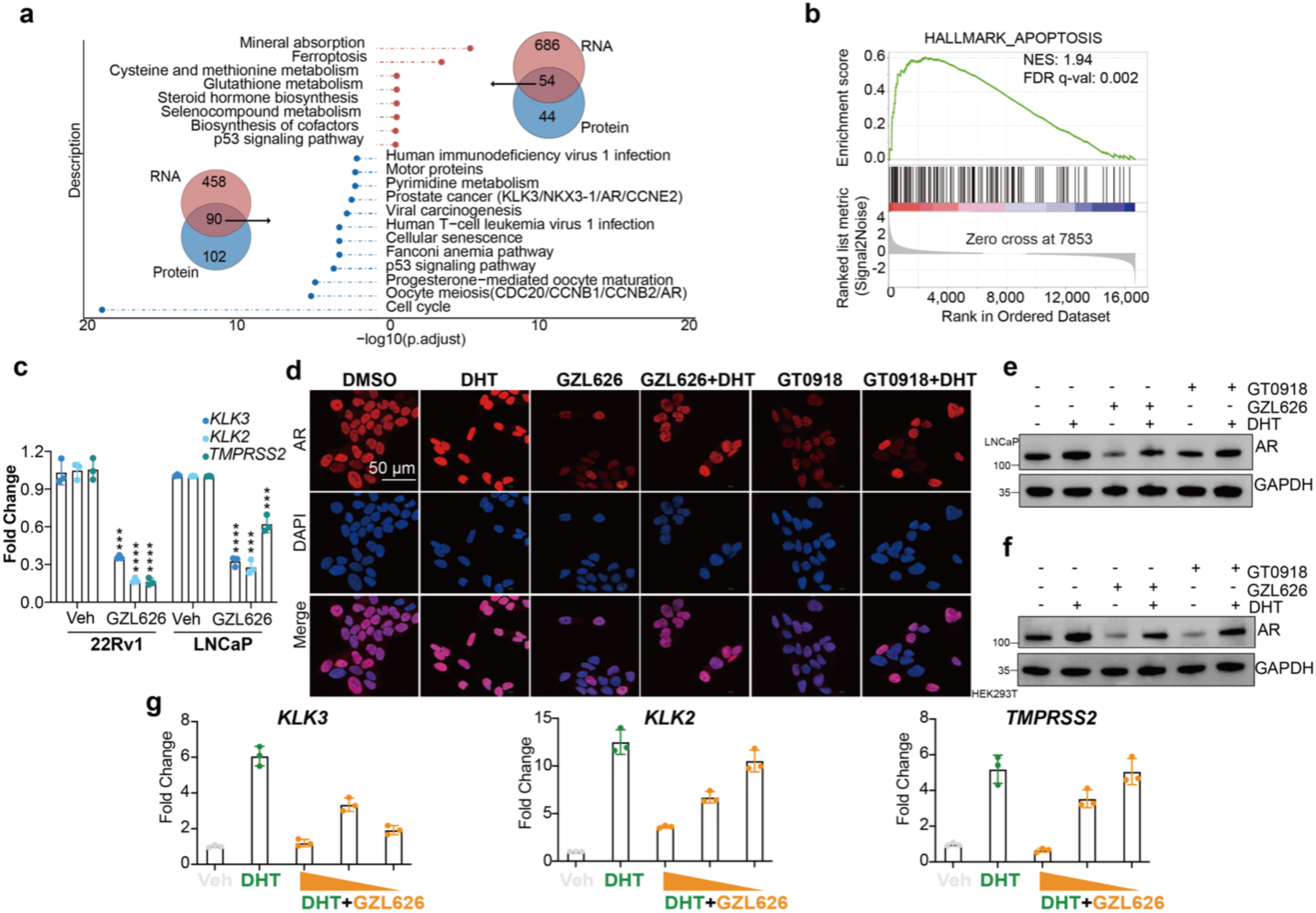
GZL626 blocks AR-dependent transcription. **a-b,** Transcriptomic and proteomic profiling of LNCaP cells treated with vehicle (Veh) or GZL626 (10 μM, 24 h, 10% CSS). (a) Venn diagram and KEGG pathway analysis. (b) Gene set enrichment for apoptosis. **c,** qPCR analysis of gene expression in LNCaP and 22Rv1 cells treated with Veh or GZL626 (10 μM, 24 h, 10% CSS). **d,** AR nuclear translocation in LNCaP cells treated with Veh, DHT (100 nM), GZL626 (10 μM), DHT + GZL626, GT0918 (10 μM), or DHT + GT0918 (24 h). Nuclei stained with DAPI. **e-f,** AR expression in LNCaP and HEK293T cells under the same treatment conditions as in **d**. **g,** qPCR analysis of gene expression in LNCaP cells treated with Veh, DHT (100 nM), or DHT + GZL626 (20, 10, 1 μM, left to right). **c, g,** n = 3 independent experiments; mean ± s.d. **d,** all images are representative. All western blot data are representative of three independent measurements. Statistical significance was defined as ***p < 0.001 and ****p < 0.0001 compared to the DMSO vehicle control.

We next examined whether dihydrotestosterone (DHT) could counteract GZL626-induced degradation. Indeed, DHT effectively compensated AR degradation in both LNCaP and HEK293T cells, a protective effect also observed with GT0918 (**Fig. 2d-f**). Nevertheless, GZL626 inhibited DHT-induced transcriptional upregulation of *KLK2*, *KLK3*, and *TMPRSS2* in a dose-dependent manner (**Fig. 2g**). These findings indicate that GZL626, like GT0918, degrades both ligand-bound dimeric AR and ligand-free inactive monomeric AR and demonstrate that GZL626 suppresses AR-driven transcriptional programs, including clinically relevant targets such as TMPRSS2, which is essential for SARS-CoV-2 entry. By simultaneously inhibiting tumor-promoting and virus-supporting gene expression, GZL626 establishes a pharmacological basis for dual antitumor and antiviral activity.

### GZL626 engages a hotspot within the intrinsically disordered NTD of AR

Unlike ARV110, none of the screened molecules, including GZL626, inhibited DHT-induced AR-ΔNTD transactivation (**Fig. 1b and Extended Data Fig. 1c**). Notably, GZL626 showed stronger degradation of AR-V7 than ARV110 (**Fig. 1e**), implicating the NTD in its antagonism. Microscale thermophoresis (MST) confirmed that GZL626 bound directly to purified full-length AR (K_d_ = 1.7 μM), ∼3-fold weaker than dihydrotestosterone (DHT; K_d_ = 0.66 μM) (**Extended Data Fig. 5a, b**). Having established that GZL626 directly binds AR, we next sought to map the binding interface within the NTD.

To determine whether GZL626 specifically interacts with the AR NTD, we performed MST and BLI using purified AR NTD fragments. Both assays showed that GZL626 binds to AR 361-537 (Tau-5) (MST K_d_ = 1.28 µM; BLI K_d_ = 2.45 µM) but not to AR 90-360 (Tau-1) (**Fig. 3a, b and Extended Data Fig. 5c**), confirming that the compound specifically targets the Tau-5 subdomain. To obtain higher-resolution mapping, we performed HDX-MS on AR 361-537 in the presence of GZL626, which identified a protected peptide spanning residues 372-394 (**Extended Data Fig. 6a-c**). To further investigate this interaction at residue-level resolution, we performed NMR structural foot printing to map the binding epitope of GZL626 on ^15^N-labeled AR NTD (90-450). The residues with the most pronounced chemical shift perturbations were H256 and L257 in Tau-1, and H384, H409, G416, and E442 in Tau-5 (**Fig. 3c-e**). Given that MST and BLI showed no detectable binding of GZL626 to the Tau-1 fragment (AR 90-360), the observed CSPs in Tau-1 likely reflect indirect conformational coupling upon compound binding to Tau-5, rather than a direct binding site. Collectively, these orthogonal biophysical assays demonstrate that GZL626 specifically engages the Tau-5 subdomain of the AR NTD, providing a molecular basis for its NTD-directed mechanism of action.

**Fig. 3.**
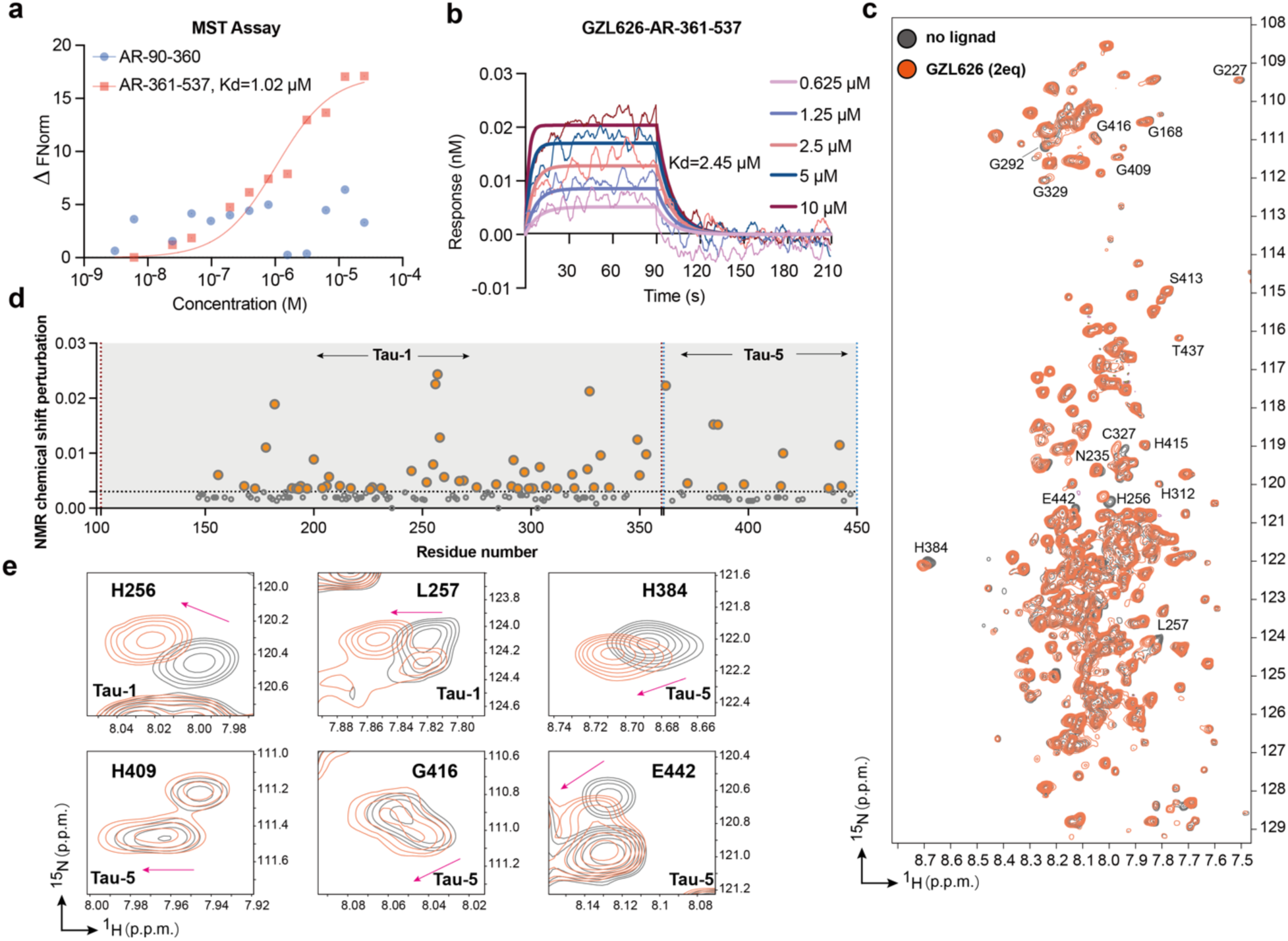
GZL626 binds the disordered domain of androgen receptor. **a**, MST binding analysis of GZL626 to purified AR NTD fragments (K_d_ values are shown in the figure). **b**, BLI binding analysis of GZL626 to AR-NTD (361-537) (the K_d_ value is shown in the figure). **c**, 2D [^1^H, ^15^N] TROSY-HSQC spectra of ^15^N labeled AR-NTD (90-450) in the absence or presence of 2 molar equivalents of GZL626, with representative residues labeled. **d**, Quantification of chemical shift perturbations (CSPs) analysis; residues with significant perturbations are indicated by dotted lines and colored circles. **e**, Selected regions of the 2D [^1^H, ^15^N] TROSY-HSQC spectra of ^15^N labeled AR-NTD (90-450) in the absence (gray) and presence (orange) of 2 molar equivalents of GZL626.

### RNF213-UBE2J2 axis mediates GZL626-induced AR degradation

To define the mechanism of GZL626-induced AR degradation, we mapped the AR interactome using proximity-dependent proteomics (APEX2 and Turbo ID) in AR-knockout HEK293T cells reconstituted with full-length AR^33^. Both N- and C-terminal tagged APEX2 fusions were degradable by AR degraders (5 μM, 12 h), whereas TurboID fusions were resistant (**Extended Data Fig. 7a, b**). We therefore selected APEX2-tagged AR for unbiased proximal labeling profiling. Following a 2-hour pretreatment with carfilzomib, cells were exposed to the vehicle or GZL626 (5 μM) for 12 hours. The cells were subsequently subjected to biotinylation, enrichment, and LC-MS analysis. (**Fig. 4a and Extended Data Fig. 7c**)^34^. GZL626 markedly reduced APEX2-AR levels, an effect attenuated by carfilzomib, indicating proteasome-dependent degradation of AR in APEX2 system (**Extended Data Fig. 7c**). APEX2 proteomics identified 108 AR-proximal proteins, including 54 known AR interactors (**Extended Data Fig. 7d-e**). UBE2J2 was identified as a novel AR-interacting protein specifically recruited in the presence of GZL626, which indicated that GZL626 may drive AR to interact with the ubiquitin-proteasome system. In UBE2J2-knockout LNCaP cells, GZL626 failed to induce AR degradation, confirming that UBE2J2 is essential for this process (**Fig. 4a-c and Extended Data Fig. 7f**).

**Fig. 4.**
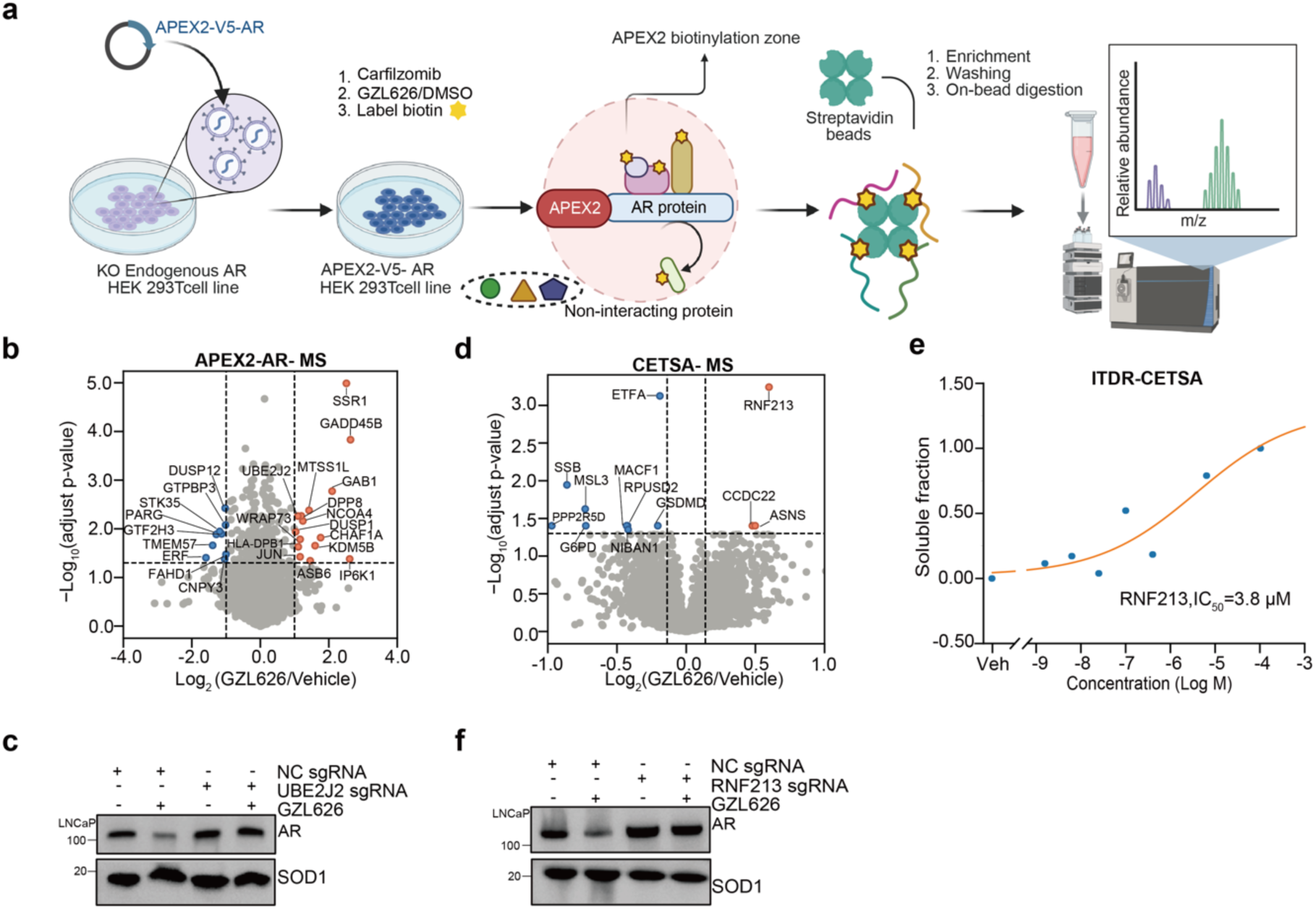
GZL626-induced degradation of AR is dependent on RNF213-UBE2J2. **a,** Schematic of APEX2-AR workflow for mass spectrometry-based identification of AR interactors. **b,** APEX2-AR proteomics in HEK293T^^AR−/−^ cells ± GZL626 (5 μM, 24 h). Statistically significant proteins (log_2_(FC) > 1; P < 0.01) are represented by blue or orange. **c,** Immunoblot of AR levels in WT and UBE2J2-KO LNCaP cells treated with GZL626 (10 μM, 24 h). **d,** CETSA-MS volcano plot of Vero E6 lysates treated with GZL626 (20 μM, 52 °C). Significant proteins (log_2_FC > 1, P < 0.01) highlighted (n = 3). **e,** ITDR-CETSA of RNF213 with GZL626 (0.01-100 μM, 52 °C). **f**, Immunoblot of AR in WT and RNF213-KO LNCaP cells treated with GZL626 (10 μM, 24 h). All western blot data are representative of three independent measurements.

To gain molecular insights into the mechanism of GZL626, we utilized cellular thermal shift assay coupled to MS (CETSA-MS) to identify the potential targets of GZL626 in Vero E6 cells. CETSA-MS revealed that E3 ubiquitin-protein ligase RNF213 was a top GZL626-engaged protein, exhibiting pronounced thermal stabilization (**Fig. 4d and Extended Data Fig. 7h**). Isothermal dose-response (ITDR)-CETSA under parallel reaction monitoring (PRM) acquisition mode to further validate the dose-dependent binding of GZL626 to RNF213 (EC_50_ = 3.8 μM; **Fig. 4e**). Owing to the low abundance of AR in Vero E6 cells, unfortunately, we were unable to identify AR in our CETSA-MS data. RNF213 contained two core functional domains: the AAA+ ATPase domain, which powered conformational remodeling through ATP hydrolysis, and the RING finger domain, responsible for mediating ubiquitin transfer^35^. RNF213 knockout abrogated GZL626-induced AR degradation in LNCaP cells, confirming functional involvement (**Fig. 4f and Extended Data Fig. 8c**). Domain analyses indicated that GZL626 does not impair RNF213 AAA+ ATPase activity, but likely interacts with its RING domain to modulate ubiquitination (**Extended Data Fig. 8 a, b**).

Biochemical reconstitution further supported a ternary mechanism. Size-exclusion chromatography (SEC) revealed co-elution of AR, UBE2J2, and RNF213, with enhanced stability in the presence of GZL626 (**Fig. 5a and Extended Data Fig. 8d**). BS3 crosslinking confirmed dose-dependent stabilization of the RNF213-AR-UBE2J2 assembly (**Extended Data Fig. 7g and Extended Data Fig. 8e**). To gain structural insights into this GZL626-stabilized assembly, we performed XL-MS in the absence or presence of GZL626 (**Fig. 5b-c**). In the DMSO-treated control, a crosslink between RNF213 K1313 and AR K846 was detected, indicative of a basal interaction interface. Upon GZL626 treatment, crosslinks involving RNF213 were broadly distributed, whereas those involving UBE2J2, and AR converged on discrete residues. In AR, intermolecular crosslinks spanned multiple domains, including the NTD (K327, K535), DBD (K609, K618), hinge region (K632, K633), and LBD (K720, K778) (**Fig. 5b**). Notably, AR K618 formed crosslinks with both RNF213 and UBE2J2 K25, suggesting that the AR DBD may serve as an important contact interface for E2-AR-E3 interactions. Intermolecular crosslinks between UBE2J2 (K17, K73, K87, K153) and RNF213 further point to extensive and integrated functional interplay between the E2 and E3 enzymes. Collectively, these results indicate that GZL626 induces a conformational rearrangement of the RNF213-AR-UBE2J2 complex into a more compact and stable assembly, consistent with the enhanced stability observed by SEC and BS3 crosslinking (**Fig. 5a-c**). Furthermore, our in vitro ubiquitination assay confirmed that GZL626 promotes AR polyubiquitination in a manner critically dependent on the RNF213-UBE2J2 complex (**Fig. 5d**). Together, these findings indicate that GZL626 facilitates the assembly of a functional RNF213-AR-UBE2J2 degradation complex, which mediates the polyubiquitination and subsequent proteasomal degradation of AR.

**Fig. 5.**
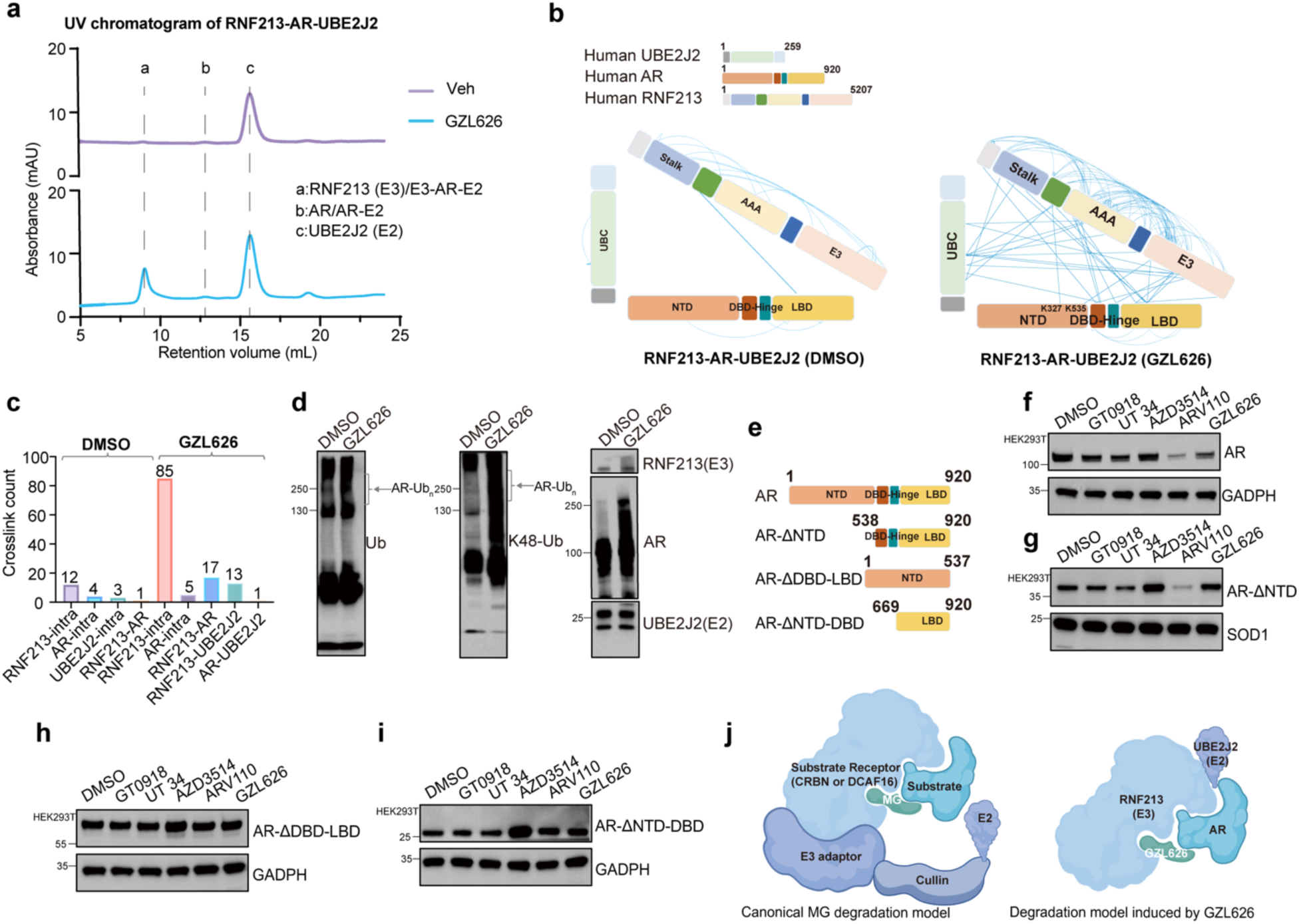
The AR-DBD as a Structural Hub Mediating GZL626-Induced AR Degradation. **a,** SEC analysis. RNF213, UBE2J2, and AR mixed at a 1:1:3 molar ratio in the absence (pink) or presence (cyan) of GZL626, run on an S200 10/300 column. **b-c,** XL-MS analysis of the RNF213– AR–UBE2J2 complex treated with DMSO or GZL626. Cross-link map showing intra-chain (arcs) and inter-chain (straight lines) interactions (b), with summary statistics of cross-link types and numbers (c). **d,** GZL626 enhances AR polyubiquitination with the involvement of RNF213-UBE2J2 complex in vitro. AR, RNF213, UBE2J2, Ubiquitin and K48-linked Ubiquitin were analyzed by immunoblotting. **e,** Schematic representation of truncated AR protein and AR mutants. **f-i,** Western blot analysis of AR levels in HEK293T cells expressing full-length or truncated AR variants, treated with vehicle or the indicated degraders (5 μM, 12 h). GAPDH and SOD1 served as loading controls. Full-length AR (AR) (f). AR lacking the N-terminal domain (AR-ΔNTD) (g). AR lacking both DBD and LBD (AR-ΔDBD-LBD) (h). AR lacking both NTD and DBD (AR-ΔNTD-DBD) (i). **j,** Schematic model comparing the canonical degradation induced by traditional monovalent glues and the degradation induced by GZL626. All western blot data are representative of three independent measurements.

Functional mapping with AR truncations further supported this model: ARV110 degraded full-length AR and AR-ΔΝTD but not AR-V7, AR-ΔDBD-LBD, or AR-LBD alone, whereas GZL626 degraded both AR and AR-V7 yet failed on isolated NTD (AR-ΔΝTD) or DBD-LBD (AR-ΔΝTD) variants, highlighting the essential role of the DBD, which might serve as a hub mediating the interaction between RNF213 and UBE2J2 (**Fig. 1e and Fig. 5e-i**). Together, these findings demonstrate that GZL626 stabilizes an RNF213-AR-UBE2J2 ternary complex, positioning the AR-DBD as a structural hub and driving proteasome-dependent AR degradation. This noncanonical E3-E2-AR underlies the antitumor and antiviral activities of GZL626 (**Fig. 5j**).

## Discussion

The activation function of AR primarily relies on the ligand-independent AF-1 region in the NTD, which contains two major transcriptional activation units: Tau-1 (102-371) and Tau-5 (361-537)^10^. In this study, we identified GZL626 as a novel AR NTD-targeting degrader. Using orthogonal biophysical approaches (MST, BLI, XL-MS, HDX-MS, and protein NMR; **Fig. 3a–e**, **Extended Data Fig. 5c and Extended Data Fig. 6a–c**), we demonstrated that GZL626 directly engages the Tau-5 subdomain of the AR NTD, with no detectable binding to Tau-1. NMR analysis revealed that GZL626 induced significant chemical shift perturbations in both Tau-1 and Tau-5. However, given the lack of direct binding to Tau-1 by MST and BLI, the CSPs observed in Tau-1 likely reflect indirect conformational coupling upon compound binding to Tau-5, rather than a direct interaction. Structurally, GZL626 and its active analogs share a distinctive core scaffold (**Extended Data Fig. 3**) that is markedly different from existing AR-targeting molecules, distinguishing them from the EPI series (non-degrading antagonists), UT-family degraders (Tau-1-targeting), and CRBN-recruiting PROTACs (e.g., BWA522, ITRI-148)^14,29,36,37^. Collectively, the distinctive scaffold of GZL626 and its analogs defines a new chemical class of AR NTD-targeting degraders, providing a novel structural framework for future drug discovery.

Mechanistically, GZL626 functions as a bivalent molecular glue degrader, with binding affinities of 1.7 µM for AR and 3.8 µM for RNF213 (**Extended Data Fig. 5a and Fig. 4e**). Integrative XL-MS analysis revealed that, in the presence of GZL626, two crosslinking sites (K327 and K535) were identified within the AR NTD and precisely mapped to the Tau-5 subdomain (**Fig. 5b**), consistent with the binding interface defined by orthogonal biophysical assays (MST, BLI, HDX-MS, and NMR). In addition, XL-MS revealed direct contacts between the AR DBD and both the RNF213 RING and UBE2J2 UBC domains, positioning the AR DBD as a central interaction hub within the ternary complex (**Fig. 5b**). Domain mapping studies functionally validated this structural model: GZL626 efficiently degraded full-length AR and AR-V7 but failed to degrade isolated NTD or DBD-LBD variants, confirming the DBD as an essential scaffold for RNF213-UBE2J2-mediated degradation (**Fig. 1e and Fig. 5e-i**).This observation suggests that GZL626 engagement of the NTD induces allosteric conformational changes that are transmitted to the DBD, a mechanism consistent with documented interdomain allostery in AR, including intramolecular NTD-LBD (N/C) interactions and NTD-dependent modulation of DNA-binding activity^3,38^. Given the high conservation of the DBD across steroid hormone receptors, this allostery-dependent degradation mode may be broadly applicable to other nuclear receptors.

AR-V7 drives resistance to enzalutamide and abiraterone in CRPC^39^. Given the limited clinical success of prior NTD-targeted candidates (e.g., EPI, UT-series).^12–14,40^. GZL626 offers a distinct clinical profile. Its degradation potency (DC_50_ ≈ 2-5 μM) is comparable to UT-series degraders but weaker than PROTACs such as ARV-110, which achieves nM-range degradation of full-length AR but is inactive against AR-V7 (**Fig. 1e, f, and Extended Data Fig. 1e, g**); nevertheless, this level of potency is largely sufficient to suppress AR-V7- and LBD mutation-driven transcription in CRPC. Compared with PROTACs, GZL626 offers several potential advantages. In principle, its lower molecular weight (<500 Da) should favor oral bioavailability and permeability^23^, although we acknowledge that the current pharmacokinetic properties of GZL626 are suboptimal, a limitation not uncommon for newly identified molecular glue leads, and further medicinal chemistry optimization will be required. GZL626 engages the AR NTD with minimal cross-reactivity to other nuclear receptors (PPARγ, ER, GR) and shows AR-dependent cellular activity, whereas ARV-110 reduces viability in AR-negative cells (**Fig. 1e, h-j and Extended Data Fig. 2a-d**). Moreover, GZL626 operates via a non-CRBN-dependent mechanism, avoiding the neo-substrate degradation (e.g., GSPT1, CK1α) associated with CRBN-recruiting PROTACs^23,32^. Collectively, GZL626 is a molecular glue degrader with an unconventional mechanism that effectively targets both full-length AR and AR-V7. Despite its moderate potency, its differentiated selectivity, avoidance of common PROTAC-related off-target effects, and activity against therapy-resistant AR variants position it as a valuable lead for further optimization in CRPC.

## Methods

### Protein expression and purification

pET-45b-His_6_-HRV3C fusions of UBE2J2 (1-226) and AR fragments (9-360, 361-537, and 447–537) were expressed in *E. coli* BL21(DE3). Cells were grown to an OD_600_ of 0.6-0.8, and protein expression was induced with 0.4 mM IPTG. The induced cells were subsequently grown overnight at 16 °C. All proteins were purified using Ni-NTA resin, concentrated to approximately 5 mg mL^-1^, and stored in a final buffer containing 50 mM HEPES (pH 7.4) and 150 mM NaCl.

For single ^15^N isotopic labeling, *E. coli* DE3 cells expressing pET-45b-His_6_-HRV3C-AR (amino acids 90-450) were grown in M9 medium (34 g/L Na_2_HPO_4_, 15 g/L Na_2_HPO_4_, 2.5 g/L NaCl) supplemented with 1× trace element solution, 2 mM MgSO_4_, 0.4% glucose, 100 µg/mL ampicillin, 1 g/L ^15^NH_4_Cl as the sole nitrogen source, and 6 mL/L Centrum multivitamins solution (prepared by dissolving one tablet in 20 mL of water). HRV3C-AR (amino acids 90–450) was purified using Ni-NTA resin, concentrated to approximately 75 µM, and stored in a final buffer containing 20 mM phosphate buffer, 100 mM KCl, 5 mM TCEP, 0.5 mM EDTA, and 10% D_2_O at pH 7.4.

pFast-Bac-His_6_-TEV-AR-FLAG plasmid was transformed into DH10 cells. Blue-white screening was used to isolate colonies containing recombinant baculoviral shuttle vectors (bacmids) and bacmid DNA was extracted by Pure link Hipure Plasmid miniprep Kit (Invitrogen #K210002). Bacmids were then transfected into adherent Sf9 insect cells in 6-well plates, using Cellfecti^TM^ II Reagent (Thermo Fisher #10362100). High titer baculoviral stocks were harvested by transfecting Sf9 suspension cultures. His_6_-TEV-AR-FLAG protein was expressed in Sf9 insect cells at 27 °C. At 24 hours post-infection, the cells were treated with 1 μM DHT and incubated for an additional 24 hours, followed by harvesting. Cells were pelleted down at 5,000 rpm for 10 minutes at 4°C, and the supernatant was removed. To lyse cells, the pellets were resuspended in 100 mL total volume in lysis buffer (50 mM Tris-HCl, pH 8.0; 150 mM NaCl; 0.5% NP40; 5% glycerol), containing 100 μL of 10 mg/mL DNase I (GOLDBIO #9003-98-9) and 1 cOmplete Tablets EDTA-free (Roche #4693132001). The cell suspension was incubated for 30 min and then sonicated for 5 min in 10-s on with 20-s off time at 30% amplitude using an Ultrasonic Homogenizer with a 900 W probe sonicator (SCIENTZ #JY92-IIDN). The resulting mixture was centrifuged at 40,000 × g for 60 min at 4 °C. The supernatant was filtered through a 0.45 μm filter unit (PALL #AVFP02SC) and incubated with 2 ml NTA affinity resin (Cytiva #GE17-5318-02) on a roller at 4 °C for 1 h. After washing the beads with 40 mL wash buffer (50 mM Tris-HCl, pH 8.0; 150 mM NaCl; 0.05% NP40; 25 mM imidazole; 5% glycerol), His_6_-TEV-AR-FLAG protein was eluted with elution buffer (50 mM Tris-HCl, pH8.0; 150 mM NaCl; 0.05% NP40; 300 mM imidazole). Eluted protein was further loaded on a Superdex 200 Increase 3.2/300 GL column (Cytiva #28990946) pre-equilibrated with gel filtration buffer (50 mM Tris-HCl, pH 8.0; 150 mM NaCl; 0.05% NP40; 5% glycerol). The purified proteins were concentrated to a concentration of ∼1 mg ml^-1^, aliquoted, flash frozen in liquid nitrogen then stored at −80 °C.

pFast-Bac-2×Strep-His_6_-TEV-RNF213-FLAG was transformed into DH10 cells. Blue-white screening was used to isolate colonies containing recombinant baculoviral shuttle vectors (bacmids), and Bacmid DNA was extracted by Pure link Hipure Plasmid miniprep Kit (Invitrogen #K210002). Bacmids were then transfected into adherent Sf9 insect cells in 6-well plates, using Cellfectin^TM^ II Reagent (Thermo Fisher #10362100). High titer baculoviral stocks were harvested by transfecting Sf9 suspension cultures. 2×Strep-His_6_-TEV-RNF213-FLAG protein was expressed at 21℃ in High Five insect cells for 3 days. Cells were pelleted down at 5,000 rpm for 10 minutes at 4°C and removed the supernatant. To lyse cells, the pellets were resuspended to 100 mL of lysis buffer (30 mM HEPES, 100 mM NaCl, 10 mM MgCl_2_, 0.5 mM TCEP, pH 7.6), containing 100 μL DNases (GOLDBIO #9003-98-9), and 2 tablets of EDTA-free protease inhibitor tablets (Roche #4693132001). The cell suspension was incubated for 30 min and then sonicated for 5 min in 10-s on and 20-s off time at 30% amplitude using an ultrasonic homogenizer with a 900 W probe sonicator (SCIENTZ #JY92-IIDN). The resulting mixture was centrifuged at 40,000 × g for 60 min at 4 °C. The supernatant was filtered through a 0.45 μm filter unit (PALL #AVFP02SC) and incubated with 2 mL anti-FLAG affinity beads (Smart-Lifesciences #SA042025) on a roller at 4 °C for 1 h. After washing the beads with 40 mL lysis buffer, 2×Strep-His_6_-TEV-RNF213-FLAG protein was eluted with lysis buffer supplemented with 250 μg/mL 3× FLAG peptide, pH 7.4. The purified proteins were concentrated to a concentration of ∼1 mg ml^-1^, aliquoted, flash frozen in liquid nitrogen then stored at −80 °C.

### NMR experiments

Two-dimensional [^1^H,^15^N]-TROSY-HSQC NMR experiments (Bruker pulse sequence = trosyf3gpphsi19) of 75 µM ^15^N-labeled AR-NTD (90-450) were performed at 278 K on a Bruker Avance III 600 MHz spectrometer equipped with TCI cryoprobes. All spectra were processed using TopSpin (Bruker, v3.6.3) and analyzed using CcpNmr Analysis Assign (v3.3.2.3) for peak picking and chemical shift assignment^41,42^. The assignment of AR-NTD (90–450) was transferred from the assignments of a smaller AR fragment 142-448 (BMRB ID: 51479). To study the binding of GZL626 to AR-NTD (90-450), we compared the 2D [^1^H,^15^N]-TROSY-HSQC spectra in the absence and presence of 150 µM compound (1:2 molar ratio). The bound sample was prepared by adding 2.5 µL of a 20 mM stock solution of GZL626 in 100% DMSO-d_6_ to the protein sample. All samples contained 20 mM phosphate buffer, 100 mM KCl, 5 mM TCEP, 0.5 mM EDTA, and 10% D_2_O at pH 7.4.

### AR degradation assays

Wild-type 22Rv1 cells and LNCaP cells (either wild-type or with knockout of RNF213 or UBE2J2) were seeded and allowed to adhere for 48 hours. The cells were treated with test compounds at the indicated concentrations for another 24 hours.

Knockout of endogenous AR was generated in HEK293T cells, which were subsequently engineered to express wild-type AR, AR mutants (H875Y and F877L), or AR truncations (AR ΔNTD, AR ΔDBD-LBD, and AR ΔNTD-DBD). These cells were seeded and allowed to adhere for 4 hours. Cells were then treated with test compounds at the indicated concentrations and incubated for an additional 12 hours.

AR protein levels were assessed by Western blotting or using the AlphaLISA, SureFire Ultra HV assay kit (Revvity, Cat. #ALSU-TANDR-A100) according to the manufacturer’s instructions.

### Rescue Experiments

For Carfilzomib and MLN4924 rescue studies, 22Rv1 cells were seeded and allowed to adhere for 48 hours, pretreated with GZL626 (10 μM) or DMSO (vehicle) for 4 h, followed by the addition of carfilzomib (10 μM) or MLN4924 (10 μM). Harvesting and Western blot analysis were performed 12 h after GZL626 treatment.

For DHT rescue studies, the cells were seeded and allowed to adhere for either 48 hours (LNCaP) or 4 hours (HEK293T), pretreated with DHT (100 nM) or DMSO for 2 h, followed by the addition of GZL626 (10 μM) or GT0918 (10 μM). Harvesting and Western blot analysis were performed 24 h after GZL626/GT0918 treatment.

### Western Blotting Analysis

Following treatment, the cells were lysed using RIPA lysis buffer (Proteintech #PR20035) supplemented with protease inhibitor cocktail (Selleck #B14001). The lysates were then centrifuged at 20,000 ×g for 20 min at 4 °C. The resulting supernatants were subjected to SDS-PAGE on 4-12% gradient gels (ACE #PR20035) and subsequently transferred to PVDF membranes (Millipore #ISEQ00010). The membranes were blocked with 5% bovine serum albumin for 1 h at room temperature and then incubated with specific primary antibodies overnight at 4 °C with gentle shaking. After washing the membranes three times with TBST (10 min per wash), membranes were probed with corresponding HRP-conjugated secondary antibodies for 1 h at room temperature, followed by another series of three 10-minute TBST washes. Protein signals were detected using an ultrasensitive ECL detection kit (Proteintech #PK10003) and captured on X-ray film. The following primary antibodies were used: anti-AR (Abcam #ab108341), anti-SOD1 (Proteintech #10269-1-AP), and HRP-conjugated anti-GAPDH (Proteintech #HRP-60004) served as loading controls.

### ARE luciferase reporter assay

The ARE-luciferase reporter plasmid contains functional androgen response elements (AREs) that drive luciferase expression upon AR binding in response to dihydrotestosterone (DHT). HEK293T cells, cultured in DMEM supplemented with charcoal-stripped FBS, were co-transfected with an AR plasmid (AR, AR ΔNTD, or AR ΔFXXLF) and the ARE-luciferase reporter. After 24 hours, cells were first treated with 20 nM DHT for 2 hours, followed by the addition of test compounds or DMSO for an additional 12-hour incubation. Luciferase activity in cell lysates was then measured using the Britelite Plus luciferase assay kit (PerkinElmer, #6066761) on a BioTek multimode plate reader according to the manufacturer’s protocol. Data were analyzed using Microsoft Office Excel and GraphPad Prism 9.

### Cell growth assays

LNCaP, 22Rv1, and PC-3 cells were cultured in RPMI 1640 medium supplemented with 10% FBS (v/v) and 1% penicillin–streptomycin (v/v). After harvesting and washing, cells were seeded in 96-well plates at a density of 1,000 cells per well in 100 µL of medium with the appropriate serum: LNCaP cells were plated in medium containing 10% charcoal-stripped FBS (csFBS), whereas 22Rv1 and PC-3 cells were seeded in medium with 10% FBS. Following a 48-hour attachment period for LNCaP and 22Rv1 cells, and a 24-hour period for PC-3 cells, compound treatments were initiated. LNCaP cells were treated with specified concentrations of GZL626 and ARV110 in 100 µL of medium containing 10% csFBS and 0.1 nM DHT for 6 days. 22Rv1 and PC3 cells were treated with the test compounds (DMSO as vehicle control) in 100 µL of 10% FBS medium for 6 days. Cell viability was assessed using the Cell Titer-Glo Luminescent Cell Viability Assay (Promega #G9242). Reagent was added to each well, and plates were shaken for 2 minutes to lyse cells, followed by a 10-minute incubation at room temperature. Luminescence was measured using a BioTek multimode plate reader. For high-content live-cell imaging, LNCaP cells were treated under the same conditions as in the viability assay and monitored in real time for 6 days using a label-free high-content imaging system (Phasefocus). All data were analyzed and plotted using GraphPad Prism 9.

### In Vitro Antiviral Activity Assay

Vero E6 cells were seeded in 96-well plates at a density of 2 × 10^4^cells per well and incubated for 24 h. To assess antiviral activity, both single concentrations and serially diluted compounds were pre-mixed with SARS-CoV-2 at an MOI of 0.01. Then, 200 µL of each mixture was inoculated onto the Vero E6 monolayer. After 72 h of infection, virus-induced cytopathic effect (CPE) was quantified using a Celigo Image Cytometer. The percentage of CPE inhibition was used to evaluate compound activity, and the half-maximal effective concentration (EC_50_) was derived from the multi-concentration data.

### Proteomic screen

LNCaP cells (800,000 cells/well) were seeded in PRMI 1640 media with 10% charcoal stripped FBS (Capricorn scientific #FBS-CS-12A) and 1% penicillin-streptomycin (Gibco #15140122) in a 6 cm dish pre-treated with 0.1 mg/mL poly-L-Lysine (Procell #PB180523) for 2 h at 37 °C. After 2 days, cells were treated with 10 µM GZL626/DMSO for 24 hours. Cell lysates were analyzed by label-free LC-MS/MS methodology.

### RNA sequencing

LNCaP cells (800,000 cells/well) were seeded in PRMI 1640 media with 10% charcoal stripped FBS (capricorn scientific #FBS-CS-12A) and 1% penicillin-streptomycin (Gibco #15140122) in 6 cm dish pre-treated with 0.1 mg/mL poly-L-Lysine (Procell #PB180523) for 2 h at 37 °C. After 2 days, cells were treated with 10 µM GZL626/DMSO for 24 hours. Total RNA of three biological replicates was extracted from the treated cells using RNAiso Plus Reagent (TaKaRa #9108) and then used for the construction of cDNA libraries by an Illumina TruseqTM RNA sample prep Kit. Sequence-by-synthesis single reads of 150-base-length using the Hiseq2500 Truseq SBS Kit (v3-HS, Illumina) were generated on the HiSeq X system. Raw sequencing data underwent quality assessment (fastqc v0.11.9) and trimming (trimmomatic v0.39), followed by alignment to GRCh38 (STAR v2.7.11) and gene quantification (featureCounts v2.0.8). The expression count matrices were normalized using logTPM transformation. Subsequently, the differentially expressed genes were selected using Bioconductor/edgeR. The gene set enrichment analyses and enrichment-related plots were carried out with GSEA software and Bioconductor/clusterProfiler.

### RNA extraction and qRT-PCR

Pre-treated LNCaP or 22Rv1 cells were lysed and extracted using RNAiso Plus Reagent (TaKaRa #9108) according to the manufacturer’s instructions. Following RNA extraction, cDNA was synthesized via reverse transcription using the Evo M-MLV RT Mix Kit with gDNA clean reagent (Accurate Biology #AG11705). Quantitative real-time PCR was performed using the 2× SYBR Green Pro Taq HS Master Mix (Accurate Biology #AG11701) on a CFX384 Touch Real-Time PCR Detection System (Bio-Rad). The PCR protocol consisted of initial denaturation at 95 °C for 30 s, followed by 40 cycles of 95 °C for 5 s and 60 °C for 30 s. Glyceraldehyde-3-phosphate dehydrogenase (*GAPDH*) was used as the internal control, and relative gene expression was calculated using the 2^−ΔΔCT^ method. The sequences of the AR-related gene primers used in this study are listed below: *GAPDH* sense (5’-AGGTCGGTGTGAACGGATTTG-3’) and antisense (5’-TGTAGACCATGTAGTTGAGGTCA-3’), *KLK2* sense (5’-GAACCAGAGGAGTTCTTGCG-3’) and antisense (5’-CCCAGAATCACCCCCACAA-3’), *KLK3* sense (5’-TACGCCTGGGCCAGATGGTG 3’) and antisense (5’-CAAGACTACGGGCCAGGCAA-3’), *TMPRSS2* sense (5’-CGTGCAGTTCGCCTCTACGG-3’) and antisense (5’-CGGGTGCTGCCCCATACTCA-3’).

### Immunofluorescence

LNCaP cells for microscopy were seeded in 35 mm dishes with a 14 mm micro-well and treated with compounds at specified concentrations. Cells were washed with PBS and fixed with 4% paraformaldehyde for 15 min at room temperature. After five PBS washes, cells were permeabilized with 0.1% Triton X-100 for 20 min and blocked with 3% BSA for 1 h. Cells were then incubated with anti-AR antibody overnight at 4 °C, followed by three PBS washes and incubation with goat anti-rabbit IgG (H&L) (Abcam #ab150080) for 2 h at room temperature. Nuclei were stained with 0.2 µg/mL DAPI (Thermo Fisher #62248). Images were acquired using a Zeiss LSM880 laser scanning confocal microscope.

### Microscale thermophoresis (MST)

Microscale thermophoresis (MST) binding assays were conducted on a Monolith NT.115Pico instrument (NanoTemper Technologies) to characterize the interaction of AR fragments (90-360 and 361-537) and full-length AR with GZL626 or DHT. Serial dilutions of each compound were prepared in PBST (PBS supplemented with 0.02% v/v Tween-20) and incubated with 5 nM AR protein. Following incubation, MST measurements were taken at 20% excitation power and 40% MST power. Dose-response curves were analyzed and plotted with GraphPad Prism.

### Bio-Layer Interferometry (BLI)

All BLI experiments were performed on an Octet System (Sartorius) at 30°C with continuous shaking at 1000 rpm. Black 96-well plates (VWR) were used, with each well containing 200 µL of BLI buffer (PBS supplemented with 0.02% v/v Tween-20). Biotinylated protein constructs (AR-90-360, AR-361-537, and AR-474-537) were immobilized onto streptavidin (SSA) biosensors (Sartorius) at optimized concentrations to achieve a stable signal response. For analyte preparation, the small molecule GZL626 was serially diluted 2-fold in BLI buffer containing 1% DMSO to generate five concentrations ranging from 0.625 to 10 µM, and the final DMSO concentration was kept constant at 1% across all samples. Control wells (including reference sensors and buffer-only wells) also contained 1% DMSO to ensure a consistent solvent background. A double background subtraction strategy was employed to correct for non-specific binding and signal drift. This included subtraction of signals from a reference sensor (no protein immobilized) and from buffer-only wells (no analyte). All binding data were globally fitted using a 1:1 binding model in the Octet data analysis software.

### Hydrogen-deuterium exchange (HDX)

A solution-phase amide hydrogen-deuterium exchange (HDX) workflow was performed on an automated system adapted from a previously described platform^43^. In brief, 5 µL of 10 µM AR-361-537 protein, with or without a 10:1 molar equivalent of GZL626, was mixed with 20 µL of D₂O-containing PBS buffer and incubated at 4 °C for various time intervals (10, 60, 300 and 900 s). After the exchange reaction, unwanted forward or back exchange was minimized, and the protein was denatured by adding a quench solution (5 M urea, 50 mM TCEP, and 1% v/v TFA). The quench solution was added at a 1:1 ratio to the protein solution. Following digestion, peptides were trapped on a C18 trap column (Thermo Fisher) and then separated on a C18 analytical column (Thermo Fisher). The separation used a 5-minute gradient (5-50% CH_3_CN, 0.3% formic acid) at 4 °C. Eluted peptides were analyzed directly using a high-resolution mass spectrometer (Thermo Fisher). All experiments were conducted in duplicate. The *.raw files were converted to *.mgf format and searched using Mascot (Matrix Science) for peptide identification. Peptides with a Mascot score of 20 or higher were retained for HDX detection. A decoy database search was conducted to exclude false positives. Finally, the intensity-weighted average m/z (centroid) of the isotopic envelope for each peptide was calculated using HDX Workbench.

### CETSA-MS sample preparation

The MS-CETSA procedure was performed as previously described with minor modifications^44^. For the single-temperature point MS-CETSA, Vero E6 cell lysates were divided into two identical aliquots (100 μg in 20 μL each), which were treated with either 20 μM GZL626 (in 1% DMSO) or DMSO alone. For the isothermal dose-response (ITDR)-CETSA, the lysates were aliquoted and incubated with a series of GZL626 concentrations or DMSO as a control. The resulting lysates were incubated at room temperature for 10 minutes, heated at 52 °C for 3 minutes in a thermal cycler, and then cooled at 4 °C for 5 minutes. After centrifugation, the resulting supernatants were collected, and the protein concentrations were determined using the BCA assay. Equal protein aliquots (50 μg) from both DMSO- and GZL626-treated samples underwent reduction with 10 mM DTT (55°C, 30 min), followed by alkylation with 20 mM IAA in the dark (room temperature, 30 min). the samples were then transferred to 10 kDa cut-off filters (Sartorius, Germany). After three washes with 100 μL of 50 mM NH_4_HCO_3_, the proteins were digested overnight at 37 °C with 1.5 μg trypsin/Lys-C in 100 μL NH_4_HCO_3_. The resulting peptides were labeled using a TMT10-plex kit, fractionated using C18 StageTips, and analyzed by LC-MS/MS. All PRM raw data were processed with Skyline (v20.2.0.286) to generate XIC and perform peak integration. Data that met the following four criteria: mass difference within ±20 ppm and dot-product (dotp) score greater than 0.7 were accepted for further analysis. For the PRM data, sigmoidal curvetting and EC_50_ calculations were performed using Python (version 3.9).

### APEX2-based proximity labeling and affinity enrichment of biotinylated proteins

HEK293T cells with endogenous AR knockout and stable expression of the APEX2-fused protein of interest were seeded in 10-cm dishes and cultured for 48 hours until reaching approximately 90% confluency. The cells were first treated with 400 nM carfilzomib (Selleck #868540-17-4) for 4 hours, followed by the addition of 5 μM GZL626 or DMSO vehicle control for the subsequent treatments. After 12 hours of GZL626 incubation, 0.5 mM alkyne-phenol reagent (MCE #1694495-59-4) was added to the culture medium for 30 minutes. Biotinylation was then induced by incubating the cells with 1 mM H_2_O_2_ for 1 minute with gentle swirling. The reaction was promptly terminated by removing the medium, and cells were washed three times with 2 mL of a quenching solution containing 10 mM L-ascorbic acid (MCE #134-03-2) and 5 mM Trolox (MCE #53188-07-1) in azide-free buffer, followed by three washes with ice-cold PBS. Cell pellets were lysed in 200 µL of EDTA-free RIPA strong lysis buffer (50 mM Tris pH 8.0, 150 mM NaCl, 0.1% SDS, 0.5% sodium deoxycholate, 1% Triton X-100) supplemented with protease inhibitor cocktail (Selleck #B14001) and super nuclease, followed by gentle pipetting and incubation at 4°C. Lysates were sonicated using a Diagenode Bioruptor Pico (30 s on/30 s off, 30 min total, 4°C) and centrifuged at 20,000 × g for 20 min at 4°C. Supernatants were collected, and protein concentrations were measured by BCA assay and normalized to 2 mg/mL. A 1 mL aliquot of lysate was subjected to click chemistry reaction using BTTAA-CuSO_4_, 2.5 mM freshly prepared sodium ascorbate (aladdin #134-03-2), and 2 mM biotin azide (Yeasen #58-85-5) for 2 hours at 37°C. Proteins were precipitated with ice-cold methyl-chloride at 4°C for 30 minutes and collected by centrifugation at 15,000 × g for 10 minutes at 4°C. Precipitates were washed twice with 1 mL of pre-cold methanol and resuspended in 100 µL RIPA lysis buffer. Approximately 2 mg of total protein was incubated with 100 µL streptavidin magnetic beads for 1 hour at room temperature with rotation. To remove nonspecific binding, beads were stringently washed as follows: twice with RIPA strong lysis buffer (2 min each), once with 2 M urea in 10 mM Tris-HCl (pH 8.0, 10 s), once with 1 M KCl in 10 mM Tris-HCl (pH 8.0, 2 min), and twice again with RIPA buffer (2 min each). For Western blotting, bound proteins were eluted by boiling in 2× loading buffer supplemented with 1 mM DTT and 5 mM biotin, separated by SDS-PAGE using 4-12% gels (ACE, #PR20035), and analyzed by immunoblotting. For proteomic analysis, streptavidin-captured proteins were subjected to on-bead tryptic digestion and prepared for mass spectrometry.

### Generation of knockout cells

Three sgRNAs per gene were designed with CHOPCHOP website for *RNF213* (5’-GTGGACCGATTTGCAGTACA-3’, 5’-CACGTGGTACCATTGCCGGA-3’ and 5’-AGTCGGTAAGAATGAACAAG-3’) and *UBE2J2* (5’-CGGATTTCCACCCGGACACG-3’, 5’-ATATGATCACTCCCAACGGG-3’ and 5’-CGACGTCTCTATACTGCCCA-3’). lentiCRISPRv2 can be digested using *BsmB*I, and a pair of annealed oligos can be cloned into digested vector. Lentivirus was produced by co-transfecting HEK293T cells with the lentiCRISPRv2 plasmid of interest together with the packaging plasmids pVSVg (Addgene #8454) and psPAX2 (Addgene #12260). The supernatant collected from transfected HEK293T cells was used to infect LNCaP cells, which were subsequently selected with 2 µg/mL puromycin. After 7 days of selection, the cells were trypsinized and reseeded into 96-well plates at a density of 0.8 cells per well to facilitate clonal outgrowth. Following 21 days of culture, individual colonies were isolated, expanded, and screened for the intended gene knockout by Western blotting or genomic DNA sequencing. Native control sgRNA was used as a reference for comparison.

### Size-exclusion chromatography (SEC) analysis

RNF213, AR, UBE2J2 (residues 1-226), and either GZL626 or DMSO (as a vehicle control) were combined in a buffer containing 20 mM HEPES and 150 mM NaCl (pH 7.4). The mixture was incubated on ice for 60 minutes. For the experiments shown in Fig. 5a and 5b, the final concentrations in the 250 µL reaction were as follows: 1 µM RNF213, 1 µM AR, 3 µM UBE2J2, and 5 µM GZL626 or an equivalent volume of DMSO. Following incubation, the samples were fractionated by size-exclusion chromatography using a Superdex 200 Increase 10/300 GL column pre-equilibrated with running buffer (20 mM HEPES, 150 mM NaCl, 1 mM TCEP, pH 7.4). The resulting fractions were collected for subsequent biochemical analyses, including SDS-PAGE and cross-linking mass spectrometry (XL-MS).

### Cross-linking mass spectrometry (XL-MS)

Cross-linking conditions were optimized using a mixture of RNF213, AR, and UBE2J2 (each at 0.25 µM) in the presence of GZL626 (0.5 µM). After a 1-hour pre-incubation at 4°C to allow complex formation, cross-linking was initiated by adding varying molar excesses of either BS3 or BS2G, followed by a 2-hour incubation at 4°C in a 20 µL volume. Control samples were prepared without a cross-linker. Reactions were terminated with 50 mM Tris-HCl (pH 8.0), and the products were analyzed by SDS-PAGE and visualized with Coomassie Blue R-250.

For the identification of cross-linked peptides, the SEC-purified RNF213-AR-UBE2J2 (residues 1-226) complex was incubated with an 800-fold molar excess of BS3 in a 100 µL reaction system at 4 °C for 2 hours, with or without GZL626. The reaction was quenched by adding 50 mM Tris-HCl (pH 8.0). Subsequently, the sample was subjected to proteolytic digestion followed by LC-MS/MS analysis. Digestion was initiated by adding 1 µL of trypsin (2 µg/µL) to 40 µL of protein sample and incubating at 37 °C for 12 hours. This was followed by sequential addition of 1 µL of chymotrypsin (1 µg/µL) twice, each followed by 4-hour digestion at 37 °C. LC-MS/MS analysis was performed on an Orbitrap Astral Zoom mass spectrometer coupled to a Vanquish Neo UHPLC system. Peptides were separated on a 15-cm C18 column over a 60-min gradient of 8-55% 80% acetonitrile in 0.1% formic acid. Data was acquired with data-dependent mode, including 3 experiments with different FAIMS CVs (−50v, −60v, −70v), and the cycle time was set to 0.6 sec for each CV setting. MS1 spectra were acquired at 180,000 resolutions. The MS1 scan range was 400-1400 m/z, the AGC target was 500%, and the maximum injection time was 5 ms. Precursor ions with charge states of 3-8 were isolated with the quadrupole mass filter and fragmented by higher-energy collisional dissociation (HCD) at a stepped normalized collision energy of 28, 32%. Quadrupole isolation was 1.6 m/z for ASTMS2 and the ms2 mass range was set to 150-2000, the AGC target was 500%, and the maximum injection time was 20 ms. The data were processed using Xlinkx in Proteome Discoverer (version 3.1) against a database containing the complex subunits. Search parameters included: KRFLWY was set as cleavage sites, up to 2 missed cleavages, precursor mass tolerance of 10 ppm, fragment mass tolerance of 0.02 Da, and oxidation (M) as a variable modification. The BS3 cross-linker was defined with a mass shift of +156.078 Da for cross-links. The false discovery rate (FDR) was controlled at < 1%.

### Ubiquitination assay

The in vitro ubiquitination assay was performed using a commercial kit (Enzo Life Science, #BML-UW9920-0001) according to the manufacturer’s instructions. The reaction mixture contained recombinant proteins (RNF213, AR, or UBE2J2) supplemented with 20× E1 ubiquitin-activating enzyme, 100 mM Mg-ATP, 20× biotinylated ubiquitin, and 10× reaction buffer, in the presence or absence of GZL626. To allow for complex formation, the mixture was first incubated at 4°C for 60 min, followed by a 30-min ubiquitination reaction at 37°C. Reactions were terminated by the addition of 2× non-reducing gel loading buffer and boiling at 100°C for 10 min, prior to immunoblotting analysis.

### NADH-coupled ATPase assay

The AAA+ ATPase activity of human RNF213 was measured using an NADH-coupled assay. Reactions were performed in a buffer containing 20 mM HEPES (pH 7.2), 200 mM KCl, 2 mM MgCl_2_, and 0.25 mM TCEP. The coupled enzyme system included 5 U/mL each of pyruvate kinase (MCE #9001-59-6) and lactate dehydrogenase (MCE #606-68-8), along with 500 µM phosphoenolpyruvate (MCE #4265-07-0) and 50 µM NADH (MCE #N6005-1G). These components were pre-mixed into a 4× master mix. For a 50 µL reaction, 12.5 µL of the master mix was combined with 12.5 µL of purified RNF213 (pre-diluted to 0.4 µM) and 25 µL of an ATP stock solution to achieve final concentrations of 0.1 µM protein and 2 mM ATP (Sigma-Aldrich #A2383-5G) supplemented with equimolar MgCl_2_. The reactions were pipetted into a 96-well plate (Beyotime #FULA962) and monitored at 30°C for 12 hours by measuring the decrease in NADH fluorescence over time using a Biotek Epoch 2 microplate reader (Agilent).

## Data availability

The RNA sequencing (RNA-Seq) datasets have been deposited in the Genome Sequence Archive (Genomics, Proteomics & Bioinformatics 2021) in National Genomics Data Center (Nucleic Acids Res 2022), China National Center for Bioinformation / Beijing Institute of Genomics, Chinese Academy of Sciences under accession code HRA013710 that are publicly accessible at https://ngdc.cncb.ac.cn/gsa-human. Source data are provided with this paper.

## Acknowledgements

We thank Guangzhou Laboratory Core Facility and Animal Center. This study was supported by grants from the Major Program of Guangzhou National Laboratory (GZNL2023A01008 to J.S. and H.C.; GZNL2025C02004 to J.S.; GZNL2023A02012 to J.S.), the National Natural Science Foundation of China (82170473 to J.S.; 22304036 to W.Q.), the Guangdong Natural Science Foundation (2021QN020451 to J.S.; 2021CX020227 to H.C.), and the Basic and Applied Basic Research Projects of Guangzhou Basic Research Program (2023A04J0161 to Q.Y.).

## Author contributions

K.W., Q.W., Y.L., Q.Y., Jiy.Z., C.H., Yu.L., J.C., and Z.Y. performed the cellular and biochemical assays. K.W., Q.W., Yu.L. and W.Q. performed proximity labeling/ CETSA-MS. K.W., J.Zha., Z.L., and J.Z. performed HDX and XL-MS. X.Q.; T.R., M.T., and H.C. performed the computational modeling. K.W., J.C., Yu.L., and Jiy.Z. performed cloning and purified proteins. K.W., Q.W., and J.S. performed the transcriptome and proteomic study. K.W., M.T., H.C., J.Z., X.C., and J.S. conceived the experiments and wrote the manuscript with input from all authors.

