## Supporting information for "A novel AR molecular glue degrader via bivalent engagement of the N-terminal domain and RNF213–UBE2J2 complex"

Kunzhong Wu *et al.*

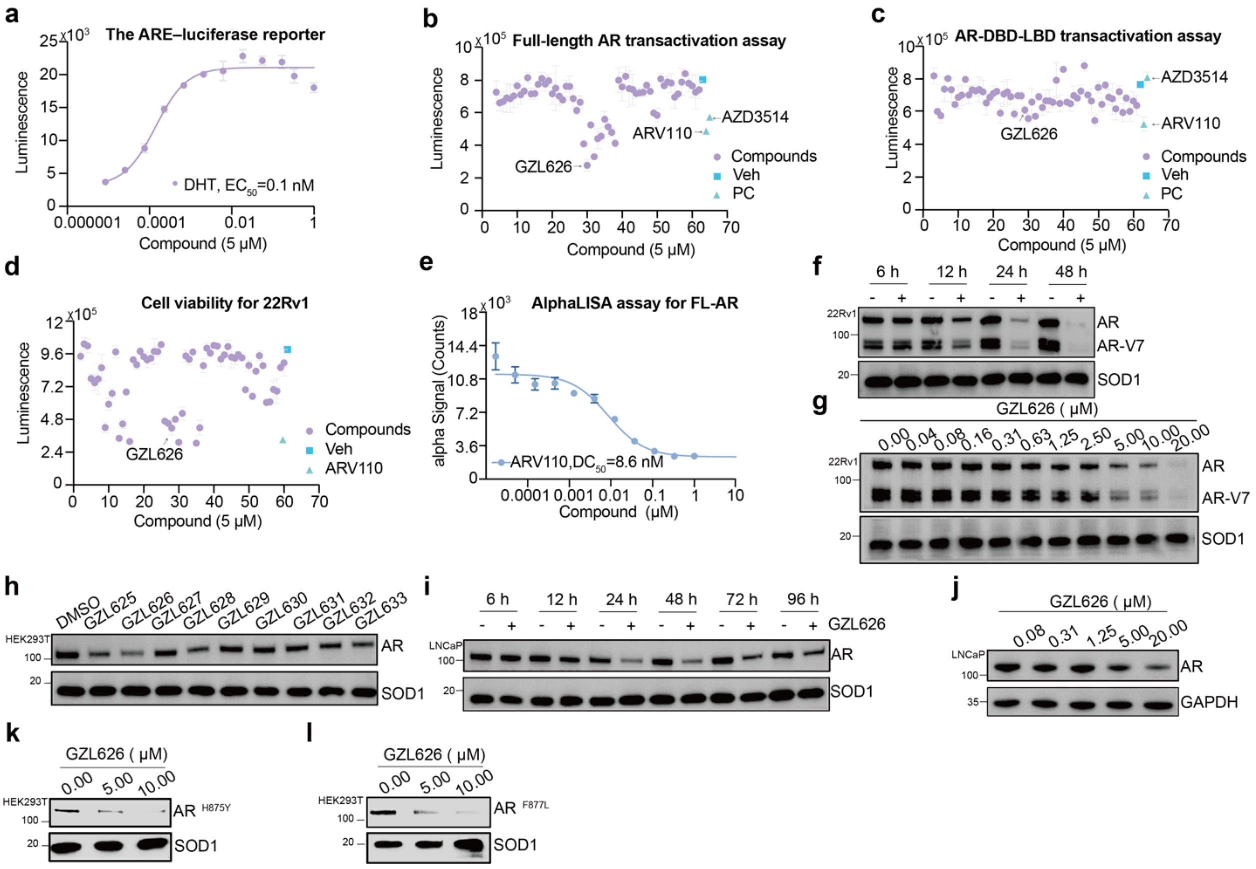


**Extended Data Fig. 1 | Discovery and characterization of GZL626.**

**a,** Dose-response curve of DHT in the ARE luciferase reporter assay ($EC_{50}$ shown). **b-c,** Effects of the in-house compound library (5 μM) on AR transcriptional activity in full-length AR (**b**) or AR lacking the N-terminal domain (AR-ΔNTD) (**c**) reporter assays with 20 nM DHT. The AR-ΔNTD construct does not represent a physiological form of AR; it was used solely as a functional tool to assess whether compound inhibition depends on the NTD. **d,** Inhibition of 22Rv1 cell proliferation by library compounds (5 μM, 6 days). **e,** Dose-dependent AR degradation in LNCaP cells treated with ARV110 (1-0.000017 μM, 3-fold dilutions, 24 h), measured by AR total detection kit. **f,** Time-dependent AR and AR-V7 degradation in 22Rv1 cells treated with vehicle or GZL626 (10 μM) for the indicated times. **g,** AR and AR-V7 levels in 22Rv1 cells treated with GZL626 (20-0.04 μM, 2-fold dilutions, 24 h). **h,** AR levels in HEK293T cells treated with GZL626 derivatives (5 μM, 12 h). **i,** Time-dependent AR degradation in LNCaP cells treated with GZL626 (10 μM). **j,** AR levels in LNCaP cells treated with GZL626 (0.078-20 μM, 24 h). **k,** AR H875Y protein levels in AR-knockout HEK293T cells overexpressing AR H875Y following treatment with GZL626 (5 or 10 μM, 24 h). **l,** AR F877L protein levels in AR-knockout HEK293T cells overexpressing AR F877L following treatment with GZL626 (5 or 10 μM, 24 h). **a-e,** n = 3 independent experiments**;** data are mean ± s.d. All western blot data are representative of two independent measurements.

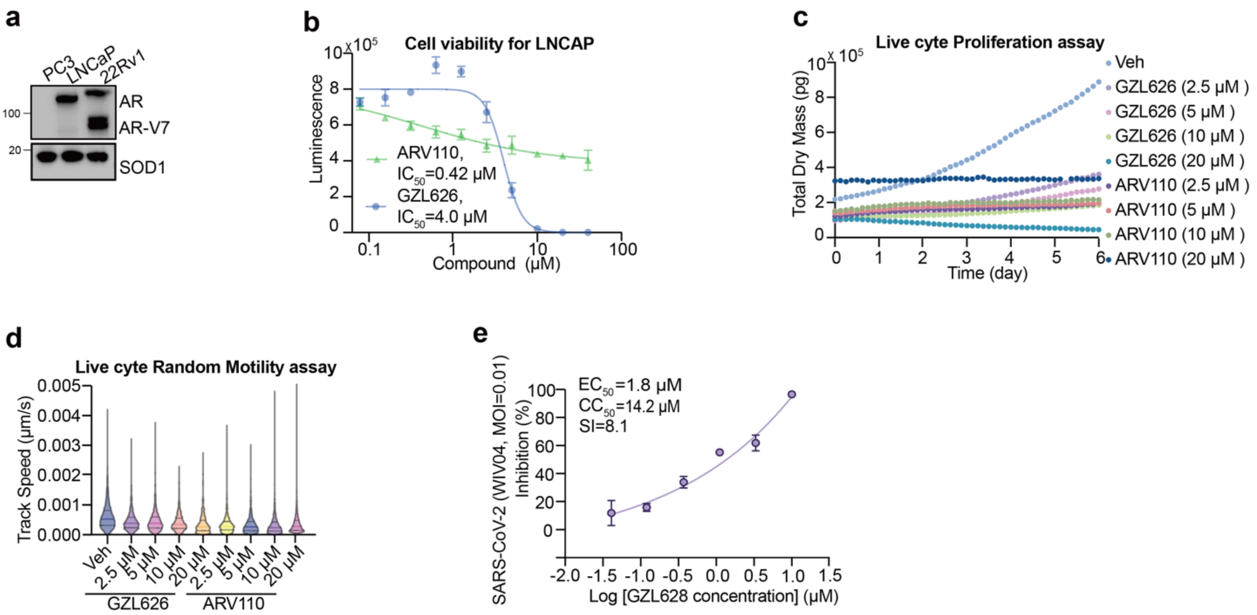


**Extended Data Fig. 2 | GZL626 suppresses LNCaP proliferation, dry mass, and motility, while GZL628 inhibits SARS-CoV-2 infection in Vero E6 cells.**

**a,** AR and AR-V7 protein levels across PC3, LNCaP, and 22Rv1 cells, assessed by western blotting. **b,** Growth inhibition of LNCaP cells treated with vehicle, GZL626, or ARV110 (40-0.08 μM, 2-fold dilutions, 6 days), with calculated $IC_{50}$ values. **c**, **d,** Effects of GZL626 (20-2.5 μM) and ARV110 on LNCaP proliferation, dry mass, and motility over 6 days, monitored in real time. **e,** Dose-dependent inhibition of SARS-CoV-2 infection by GZL628 in Vero E6 cells with $EC_{50}$ and $CC_{50}$ values shown. **b-e,** n = 3 independent experiments**;** data are mean ± s.d. All western blot data are representative of two independent measurements.

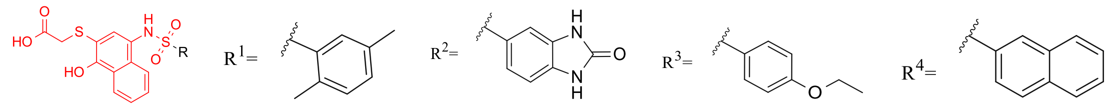

GZL625 GZL626 GZL627 GZL628

**Extended Data Fig. 3 | Chemical structures of GZL625, GZL626, GZL627 and GZL628.**

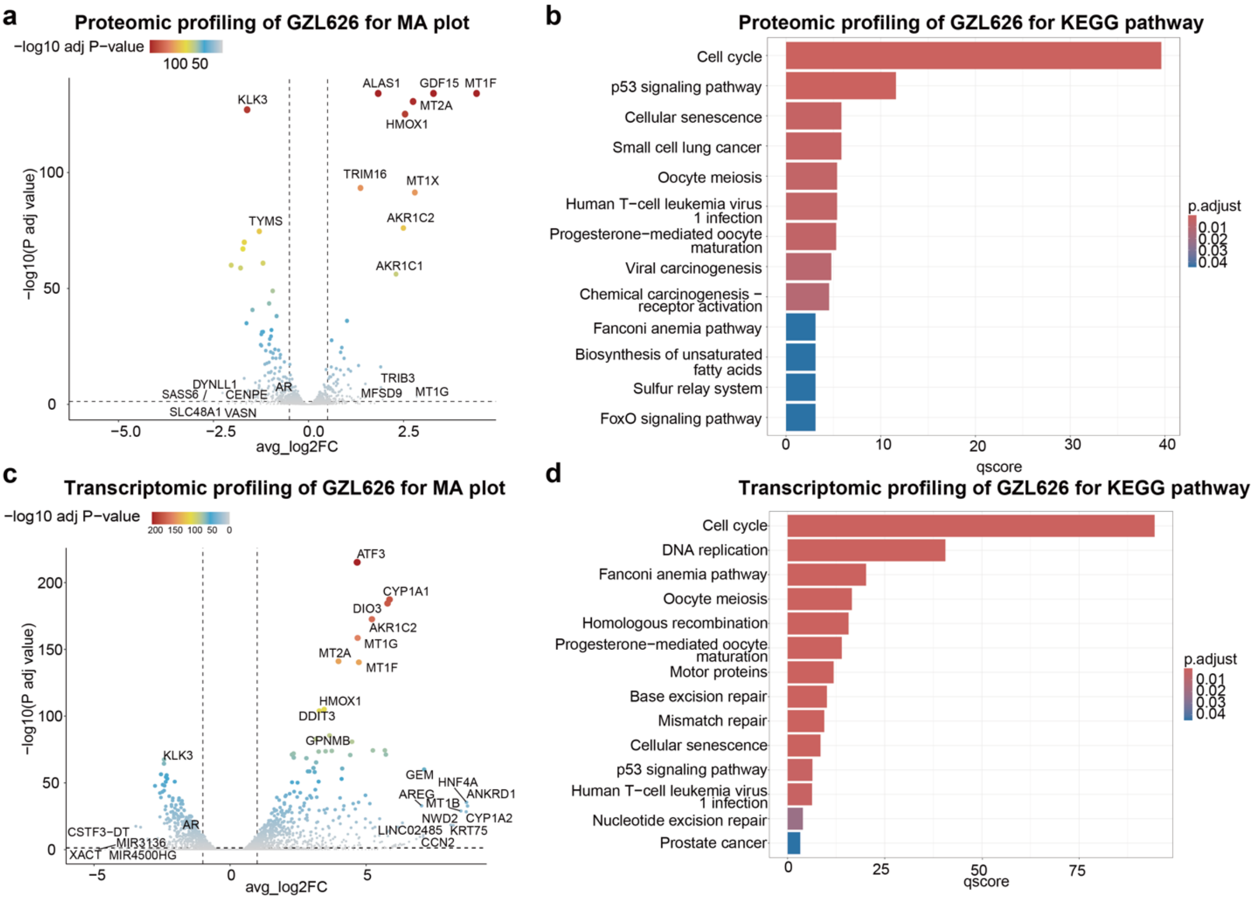


**Extended Data Fig. 4 | Transcriptomic and proteomic profiling of GZL626.**

**a-b,** Proteomic profiling of LNCaP cells treated with vehicle (Veh) or GZL626 (10 µM, 24 h, 10% CSS). (**a**) MA plot of protein abundance; significantly changed proteins ($log_2FC > 1$; $P < 0.01$) highlighted, with selected proteins annotated. (**b**) KEGG pathway enrichment of significantly altered proteins. **c-d,** Transcriptomic profiling of the same cells by RNA sequencing. (**c**) MA plot of transcript abundance; significantly altered genes ($log_2FC > 1$; $P < 0.01$) highlighted, with selected genes annotated. (**d**) KEGG pathway enrichment of significantly altered genes. **a–d:** n = 3 biologically independent replicates per group.

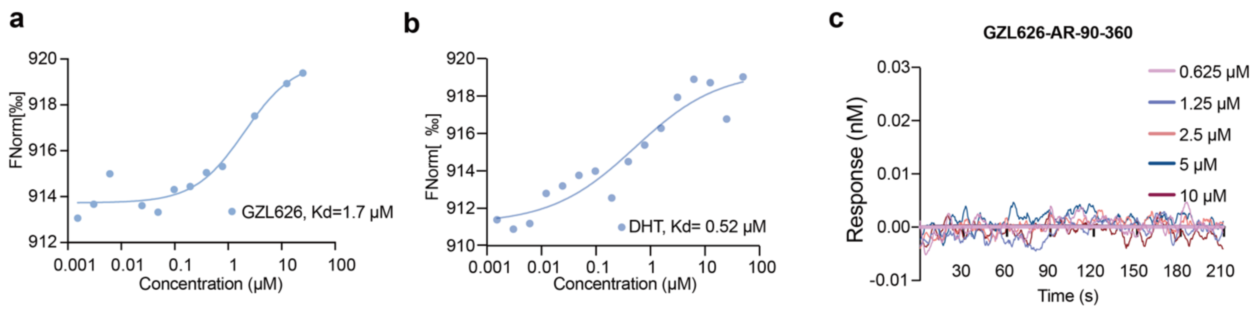


**Extended Data Fig. 5| GZL626 binds to full-length AR but not to the AR 90–360 fragment.**

**a**, Microscale thermophoresis (MST) analysis of full-length AR incubated with increasing concentrations of GZL626. $K_d$ value is shown in the figure. **b**, Microscale thermophoresis (MST) analysis of full-length AR incubated with increasing concentrations of DHT. $K_d$ value is shown in the figure. **c**, BLI binding analysis of GZL626 to the AR 90-360 fragment. No detectable binding was observed.

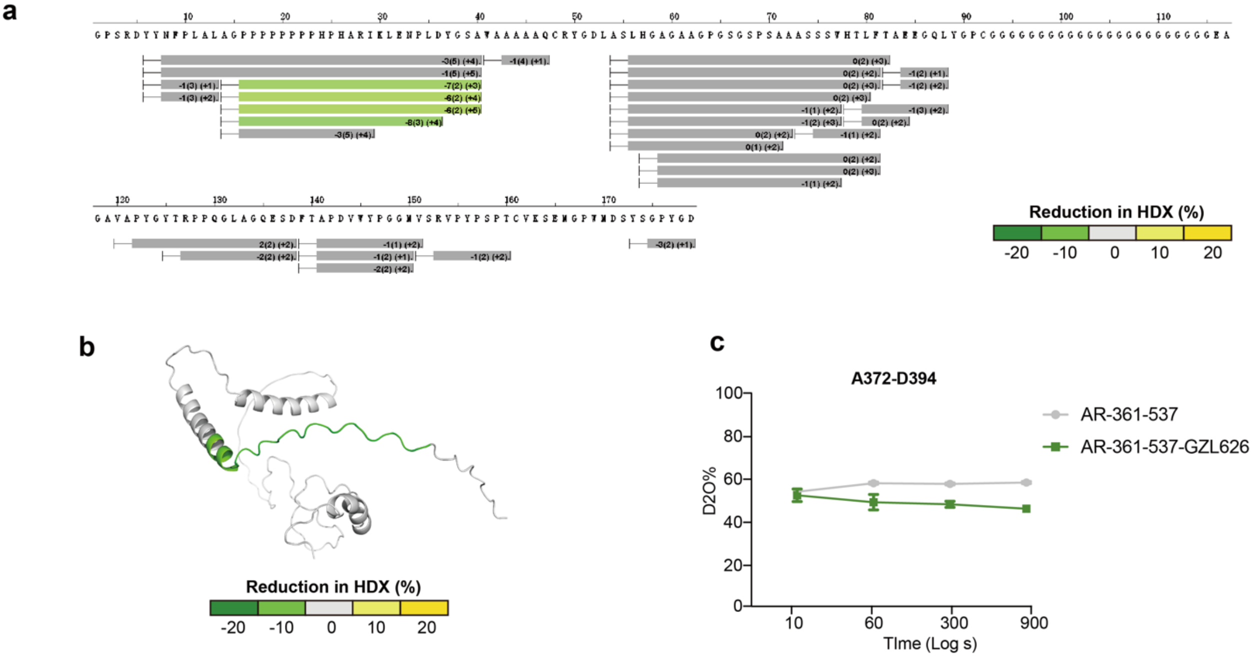


**Extended Data Fig. 6| HDX-MS analysis reveals GZL626 binding to AR-361–537.**

**a**, Percentage change in deuterium uptake (scale at the bottom) plotted against the peptides analyzed in the experiment. **b**, Differential HDX-MS analysis of AR-361–537 ± GZL626, shown as the change in deuterium uptake mapped onto the AlphaFold 3 predicted structure. **c**, Deuterium uptake plots for peptides affected by ligand binding in the absence (gray) or presence (green) of GZL626. Data represent mean ± SEM from three experimental replicates.

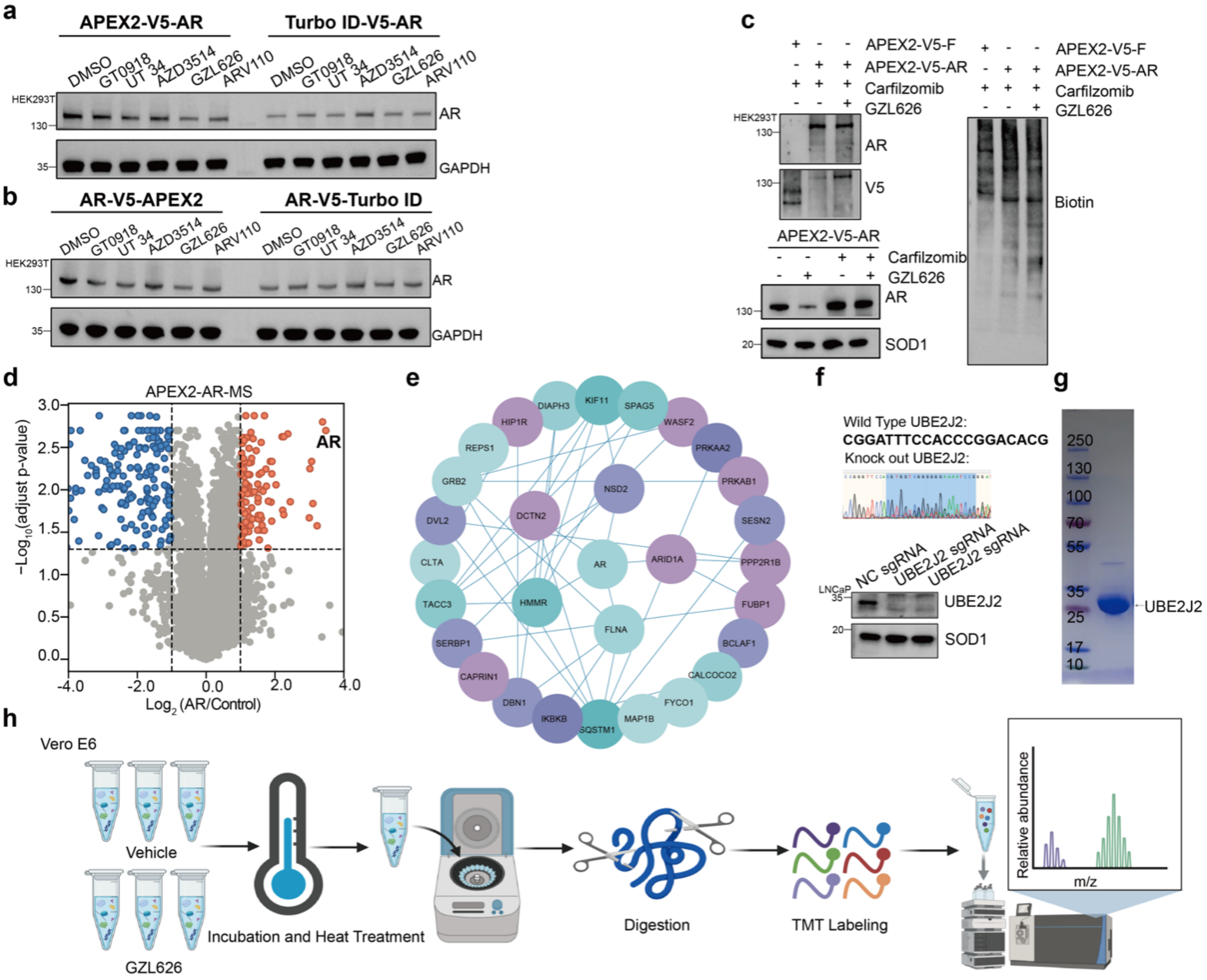


**Extended Data Fig. 7 | GZL626 recruits UBE2J2 to degrade AR.**

**a-b,** AR levels in HEK293T cells expressing APEX2-AR or TurboID-AR treated with GZL626 (5 μM, 12 h) or clinical-stage AR degraders (GT0918, UT-34, ARV110, AZD3514). AR was detected by western blotting. GADPH served as a loading control**.** c, HEK293T^$^{AR-/-}$ cells expressing APEX2-V5-AR treated with carfilzomib (0.4 μM), biotin (50 μM), ± GZL626 (5 μM). AR, V5, and biotinylated proteins (streptavidin pull-down) were analyzed by immunoblotting. SOD1 served as a loading control. **d,** Fold-change in abundance of streptavidin-enriched proteins in APEX2-V5-AR versus control, with significant proteins ($log_2FC > 1$, $P < 0.01$) highlighted (n = 3). **e,** Protein-protein interaction (PPI) network from STRING (v11.5, confidence > 0.4). Hub proteins (light blue) ranked by MCC score in Cytoscape. **f,** CRISPR-Cas9 UBE2J2-knockout cell line generation and verification. **g,** Coomassie-stained gel of purified UBE2J2 (Residues 1-226). **h,** Workflow of MS-CETSA in Vero E6 cells. All western blot data are representative of two independent measurements.

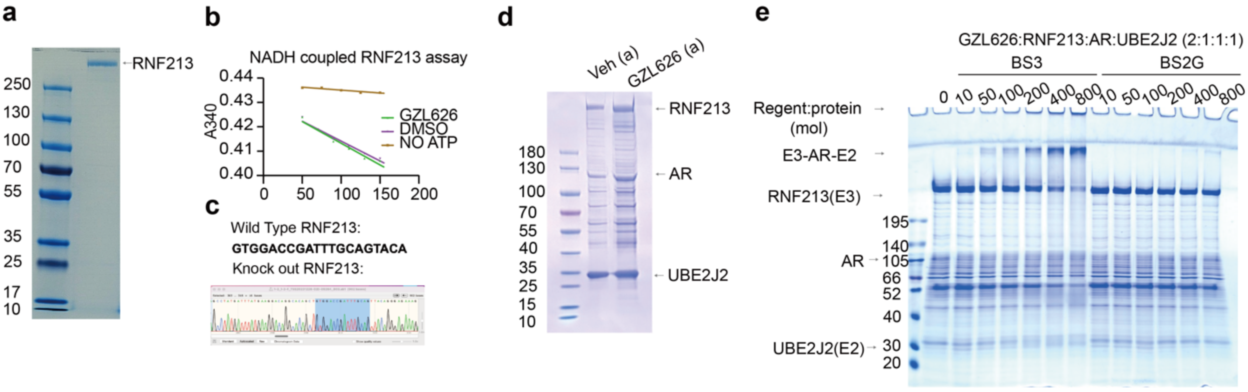


**Extended Data Fig. 8 | GZL626 induces AR degradation by promoting the formation of a functional RNF213-UBE2J2-AR complex.**

**a,** Coomassie-stained gel of purified human RNF213. **b,** ATPase assay of RNF213 treated with vehicle or GZL626 (5 μM). **c,** Generation and validation of RNF213-knockout cells using CRISPR-Cas9. **d,** Coomassie-stained gel of SEC fractions from **Fig. 5a**. **e,** Crosslinking assay showing GZL626-induced formation of an RNF213 (E3)-AR-UBE2J2 (E2) complex, visualized by Coomassie staining.
